# Convergence and conflict among telomere specialized transposons across 60 million years of Drosophilid evolution

**DOI:** 10.1101/2025.06.09.658640

**Authors:** Jae Hak Son, Matthew A. Lawlor, Mahek Virani, Weihuan Cao, Mia T Levine, Christopher E. Ellison

## Abstract

The *Drosophila* telomere is one of the best-studied examples of active transposable elements (TEs) benefitting, rather than harming, a host genome. All *Drosophila* species lack telomerase and instead have telomeres composed of head-to-tail arrays of specialized retrotransposons. These TEs ostensibly act as mutualists by elongating chromosome ends, but evidence from species closely related to *Drosophila melanogaster* suggests that telomeric transposons may also antagonize their host genome. Importantly, the limited number of *Drosophila* species characterized thus far has precluded our ability to delineate idiosyncratic from universal evolutionary forces and genetic mechanisms that shape the history of these TEs. Here, we have surveyed long-read genome assemblies of over 100 species of *Drosophila*, identifying a total of 396 telomeric TE families. Our findings show that these telomere-specialized elements evolve rapidly and also undergo striking convergent evolution: the complete loss of telomeric TEs has occurred repeatedly across the genus while individual telomeric TE lineages have repeatedly lost one of their two protein-coding genes. These elements have also repeatedly undergone horizontal transfer between distantly related *Drosophila* lineages and have repeatedly captured host gene fragments that promote their selfish suppression of host TE-silencing systems. Furthermore, telomere specialization itself appears to have evolved convergently, as some non-telomeric families have gained the ability to target their insertions to telomeres. These results provide unprecedented resolution into the evolution of these unusual TEs and highlight several novel mechanisms by which they evolve in conflict both with each other and their host genome despite the essential telomere function they provide.

## Introduction

Transposable elements (TEs) selfishly replicate in the genomes of most species, often at the expense of their host. Although most new TE insertions are either neutral or deleterious, TE copies can also be co-opted by their host genome to serve a beneficial purpose, usually as a source of new genes or novel gene regulatory elements (1–5). A large number of eukaryotic genes and regulatory sequences have been acquired from TEs. For the vast majority of these cases, the co-opted TE no longer actively mobilizes in the host genome and may evolve beyond recognition. This is the case with many TE-derived host genes where the sequence signature of their TE progenitor has eroded over evolutionary time, via deletions and other mutations, to the point where it is undetectable. Importantly, this co-option process resolves the conflict between TE and host by removing the potentially deleterious consequences of TE mobilization from the adaptive allele carried by the TE.

However, not all adaptive TEs are immobile. Recent work from a variety of species raises the possibility that active TEs can perform essential host functions, a process termed “TE addiction” (6, 7). This phenomenon refers to a period of host/TE cooperation that occurs during the transition of the TE from a selfish parasite to a fully co-opted, immobile element (7). Arguably the most well-characterized example of an active TE providing a benefit to its host genome involves the telomeric transposons of the *Drosophila* genus. All “true fly” Dipteran species lack telomerase activity (8) and the vast majority of *Drosophila* species have telomeres composed of head-to-tail arrays of specialized retrotransposons (9–11). These TEs are derived from the Jockey clade of LINE retrotransposons and were first characterized in *D. melanogaster*, where three telomeric TE families have been identified: *HeT-A*, *TAHRE*, and *TART*, collectively known as *HTT* elements (10, 12–14). The Jockey clade of LINEs generally contains two open reading frames (ORFs): a gag-like ORF1, characterized by major homology region (MHR) and zinc knuckle (CCHC) motifs, which play roles in nuclear localization and multimerization (15), and ORF2, which contains both endonuclease and reverse transcriptase domains (16).

These telomeric elements provide a clear benefit to their host: they prevent chromosome shortening due to the incomplete replication of chromosomal DNA during cell division (i.e. the end-replication problem (17)). Furthermore, the transcriptional silencing of these elements by the host genome may promote the assembly of a multi-protein complex, known as Terminin (analogous to Shelterin), that protects the chromosome ends from inappropriate double-strand break repair (18, 19). However, there is substantial evidence that these telomeric TEs are evolving in conflict with their host genome rather than existing exclusively as mutualists (20–22). For example, genes required for telomere integrity tend to evolve rapidly under positive selection (23), a signature of genetic conflict. Furthermore, telomeric TEs themselves show dynamic patterns of evolution. Recent work found multiple cases of replacement of telomeric TE lineages among closely related *Drosophila* species. Notably, the same study found that telomeric TEs were completely lost in *D. biarmipes* (24), suggesting that this ostensibly essential, mutualistic relationship between TE and host is not universally required for telomere maintenance. Furthermore, swapping into *D. melanogaster* an adaptively diverged version of a Terminin protein from a close relative resulted in a burst of transposition to telomere ends, suggesting that telomere binding proteins in *Drosophila* play a role in constraining telomeric TE activity, in addition to their end-capping function (25–27).

The piRNA pathway also constrains the activity of telomeric TEs (28–31). Many TE transcripts, including those from telomeric TEs, are cleaved into sense and antisense piRNAs via the Argonaute proteins Ago3 and Aub (32). These piRNAs then direct the formation of heterochromatin at TE loci in the genome via another Argonaute protein, Piwi, and a variety of accessory proteins including Panoramix and Nxf2 (33–36).

Mutations in the piRNA pathway result in increased retrotransposition of telomeric transposons to chromosome ends (29). Thus, telomeric TEs can act selfishly: unless actively constrained by the host genome, they hyperproliferate at chromosome ends, which has previously been shown to negatively affect host fitness (37).

Additional evidence of ongoing host-TE conflict comes from the *D. melanogaster TART* telomeric TE, which produces abundant sense and antisense piRNAs across its entire length (22). The imprecise replication of a variety of TE families, including LINE elements, has been shown to result in the acquisition of host DNA sequence by the TE, a phenomenon known as transduction or gene capture (38–40). Through this process, *TART* has captured a portion of the host piRNA pathway gene *nxf2* (22). A subset of the abundant antisense piRNAs produced from *TART* are able to target *nxf2* for suppression, which represents a form of host anti-silencing (5). The host gene *nxf2* is evolving rapidly specifically in the region that was captured by *TART*, likely due to selection to escape targeting by *TART*-derived piRNAs (22). While these results support an antagonistic relationship between telomeric TEs and their host genomes, other work raises the possibility that the rapid evolution of telomeric TEs is instead a consequence of their adaptation to the telomeric niche, which itself is inherently unstable (41, 42).

Are telomeric TEs in the process of being tamed by their host genome, as predicted by the “TE addiction” theory? Or, do they remain fundamentally selfish, despite the beneficial function they provide? If these elements are true mutualists, we would expect their evolutionary diversification to be tightly coupled to that of their host, resulting in a pattern of co-speciation, as has been observed for obligate endosymbionts of various insect species (43). Under this scenario, the previously described examples of gene capture and loss of telomeric TEs would represent rare edge cases where mutualism has reverted back to parasitism. On the other hand, a lack of co-speciation, repeated gene capture and loss of telomeric TEs would be more consistent with a parasitic relationship. These questions remain unresolved, in part because telomeric TEs have been described in only a tiny fraction of *Drosophila* species (16 (11, 24) out of more than 1,600 species in the genus (44)). Indeed, this limited picture of telomeric TE diversity precludes our ability to delineate universal from idiosyncratic evolutionary forces and genetic mechanisms that shape the history of these seemingly beneficial but active transposable elements.

Here, we manually curate 396 telomere-specialized TE families from over 100 species of *Drosophila*. Through phylogenetic and sequence analysis of these transposons, we show that these elements evolve rapidly and have undergone complete loss events on at least ten separate occasions across the genus. We identify surprising patterns of evolution, including the frequent horizontal transfer of these TEs, the convergent evolution of telomere localization, and the potential neofunctionalization of their second open reading frame, known as ORF2, whose ancestral function is conserved across all LINE transposons, including human LINE-1 elements. Finally, we show that, across the *Drosophila* phylogeny, fragments of host piRNA pathway genes have been captured repeatedly by distinct telomeric TE families. Together, these results provide strong support for antagonistic coevolution between telomere-specialized TEs and their hosts, despite the essential function they serve.

## Results

### Identification of telomere-specialized retrotransposons

We searched for non-LTR retrotransposons from the Jockey superfamily in long-read genome assemblies from a total of 106 *Drosophila* species and three outgroup species of Drosophilidae (**Table S1**). We used an automated pipeline to detect ORFs with homology to known *Jockey* superfamily ORF1 and ORF2 peptides in each assembly, followed by a phylogenetic approach to assign ORFs to telomeric versus non-telomeric clades (see Methods, **Figure S1**). We then used manual curation to generate full-length consensus sequences for each TE family and to confirm that candidate telomeric TEs form head-to-tail arrays at contig ends, consistent with telomere-specialization (see Methods, **Figure S1**, **Figure S2**).

We identified a total of 396 putatively telomere-specialized TE families across 109 species (**Table S2**). Of the 396 telomeric TE families, we detected 188 families in head-to-tail arrays at the extreme ends/termini of gene-rich, megabase-long contigs in their host species’ genome assembly. 189 TE families were found in multiple head-to-tail copies but only on short contigs lacking genes that would allow their assignment to chromosome arms. The remaining 19 TE families were present in a single copy in their host species genome assembly but were supported as telomeric based only on their position in our ORF1 and/or ORF2 gene trees (**Figure S3**). We were unable to identify telomeric TEs in a total of 15 species, including the previously reported case in *D. biarmipes* (24). In three of these species (*D. kurseongensis*, *D. orena*, and an undescribed species from the *funebris* group, labeled as *D. sp.St01m* in (45)), we identified fragments of telomeric TEs lacking intact ORFs, but no full-length telomeric TEs (**Table S3**). To confirm the absence of active telomeric TEs in all 15 species, we analyzed the raw genome sequencing reads and recovered only the same telomeric TE fragments present in the genome assemblies of the three species above, confirming the absence of telomeric TEs in these species. In total, we infer that telomere-specialized TEs were independently lost at least 10 times across the genus (**Figure 1**).

**Figure 1.**
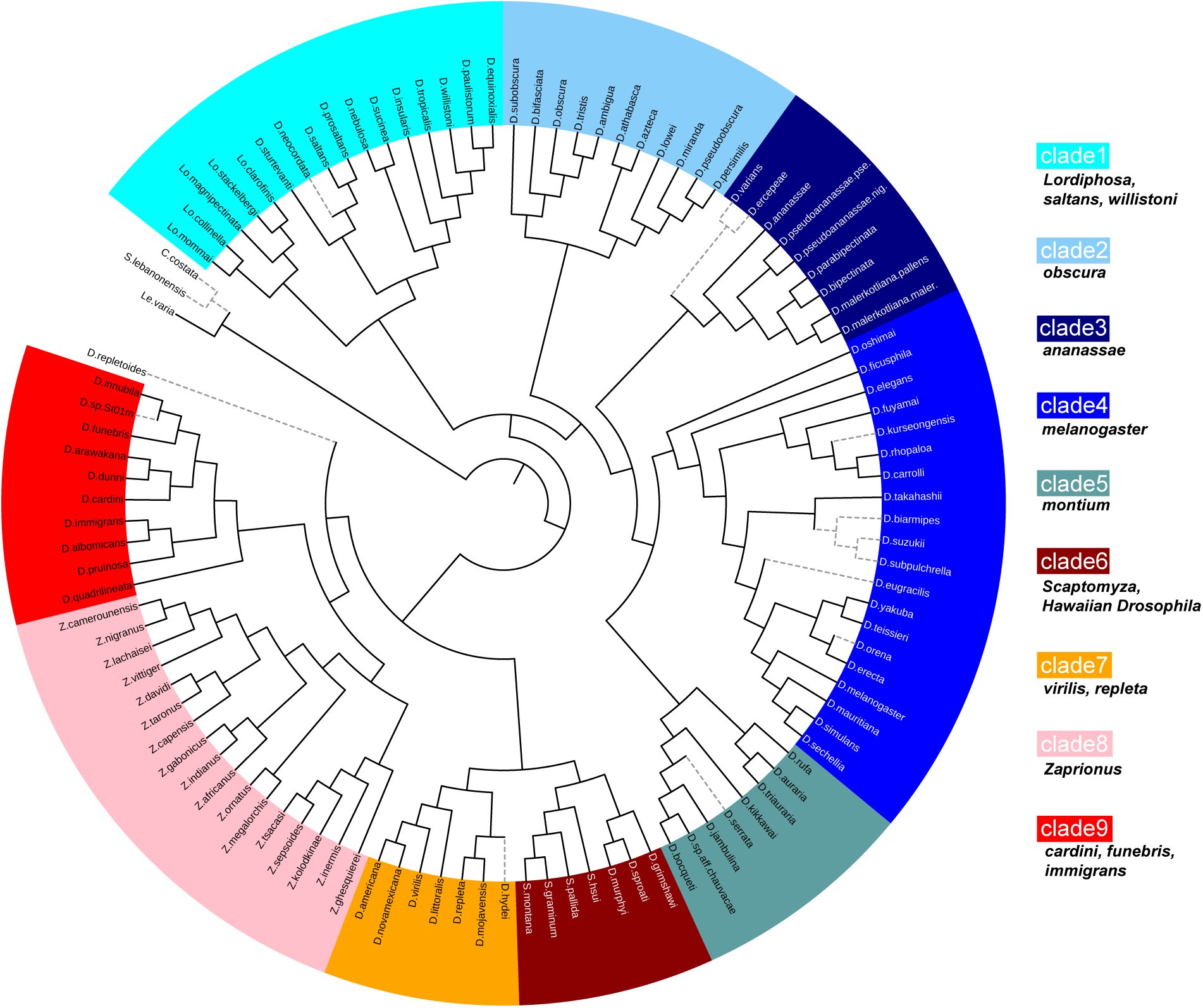
Evolutionary relationships of the species included in this study. Colors correspond to major *Drosophila* clades, as delineated in (46). The dashed grey branches indicate species for which no telomeric TEs were identified.

In the remaining species, we identified ∼4 telomere-specialized families per species, on average (**Figure S4**). On average, full-length elements were 8.00 kb in size and present in approximately 10 copies per genome, based on read depth (see Methods)(**Figure S4**). The vast majority of species with telomeric TEs have at least one telomeric TE family that carries both ORF1 and ORF2; however, we identified full-length TEs carrying only ORF1 in the genomes of *Z. ghesquierei* and *D*. *rufa* but only fragments of telomeric TEs with ORF2. Overall, ∼60% of telomeric TE families contained both ORF1 and ORF2, while ∼40% contained ORF1 only, having lost ORF2 (**Figure S4**). We also found 18 TE families where ORF2 was present but ORF1 was missing, a structure that has not been previously reported. In five of these families, ORF1 was interrupted by a premature stop codon while the remainder completely lacked ORF1 sequence (**Table S2**). These TEs have significantly reduced copy numbers compared to TE families with either ORF1 only or both ORF1 and ORF2 (Wilcoxon test *P* = 0.0004 and *P* = 0.0022, respectively)(**Figure S5**). We therefore conclude that these likely represent older fragments of previously active TEs. Taken together, these results suggest that the genomes of the vast majority of *Drosophila* species contain multiple families of telomere-specialized retrotransposons and that the dependence of ORF1-only telomeric TEs on ORF2 proteins encoded by other telomeric TE families is widespread across the genus. The two species for which we were unable to identify any intact telomeric TEs with ORF2 (in either the genome assembly or the raw reads) raise the possibility that some telomeric TEs could be using ORF2 from other non-telomeric Jockey clade TEs for replication.

### Evolutionary Diversification of telomere-specialized retrotransposons

We used ORF1 and ORF2 peptide sequences to create phylogenetic trees showing the evolutionary relationships among all 396 telomeric TE families. We identify 6 major clades of telomeric retrotransposons across 109 *Drosophila* species. These clades are present in both our ORF1 and ORF2 trees and include all original telomeric TE clades described by Villasante *et al* : *TR1*, *TR2*, *TR3*, *TR4*, and *HTT*, though we note that our more comprehensive trees suggest the *TR4* clade is actually part of the *HTT* clade (**Figure 2A, Figure S6**) (11). We therefore use *HTT* here to encompass both groups (**Figure 2A, Figure S6**). Our results confirm the monophyly of the Villasante *et al* clades and also provide evidence of two novel clades, which we name *TR5* and *TR6*. Many species contain diverse assemblages of telomeric TEs: 94 species have telomeric TEs from more than one *TR* clade, with 18 species harboring three or more *TR* clades (**Figure 2A, Table S2**).

**Figure 2.**
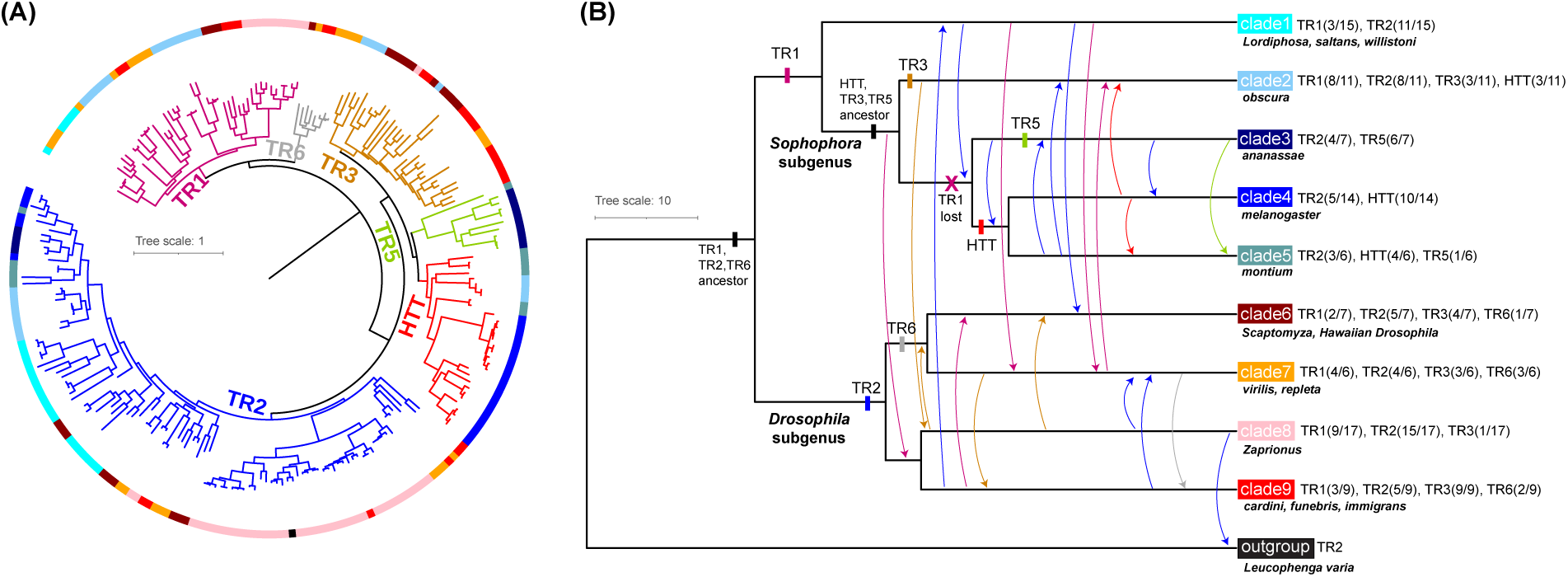
Evolutionary history of telomeric TE families. (A) Phylogeny of telomeric TE families inferred from ORF2 peptide sequences. Six major telomeric retrotransposon (*TR*) clades were identified, each with 100% bootstrap support (see **Figure S6**). Naming scheme is based on *TR* clades originally described in (11), with the addition of the novel *TR5* and *TR6* clades identified here. The tree branches are colored by *TR* clade, while the outer circle colors correspond to the host species clades shown in Panel B. The tree is rooted using ORF2 sequences from non-telomeric jockey clade TEs (see **Figure S6**). (B) Dated species tree showing major events in the diversification of *Drosophila* telomeric TEs based on species tree/TE tree reconciliation (see Methods). Arrows and rectangles are colored based on the *TR* clades shown in Panel A. *TR* clade origins are indicated by filled rectangles while losses are indicated by an “X”. Arrows show horizontal transfer events. Parentheses next to the major *Drosophila* species clades indicate the number of species harboring the corresponding *TR* clade over the total number of species within each species clade.

We used our ORF1 tree to investigate the evolutionary relationships among the abundant ORF1-only TEs found across the *Drosophila* genus. The location of these TEs within our ORF1 tree suggests that ORF2 has been repeatedly lost across dozens of independent TE lineages (**Figure S7**). However, we also find monophyletic clades containing between two and six ORF1-only TE families, which suggests that not all ORF1-only TEs arise from independent loss of ORF2 (**Figure S7**). Instead, ancestral ORF1-only TE lineages can birth new ORF1-only TE lineages. The repeated loss of ORF2 from telomeric TEs is particularly striking given that this phenomenon does not seem to occur in non-telomeric *Jockey* elements (47).

We next sought to investigate the evolutionary history of these telomeric TEs across the *Drosophila* genus. In a mutualistic relationship, where the symbiont is transmitted vertically from parent to offspring, its evolution will be tightly coupled to that of its host, resulting in a pattern of co-speciation. Such a pattern has been observed in a variety of obligate endosymbionts (43). Thus, if telomeric TEs are acting as mutualists, the major telomeric TE clades should mirror the major clades found in the *Drosophila* species tree. We would also expect that the most ancient TE clade should be found broadly, across all extant *Drosophila* species, since its origin would coincide with or predate the most recent common ancestor of the genus. Instead, we find that many distantly related species harbor closely related telomeric TEs. For example, *TR1*, *TR2*, and *TR3* clade elements are found in species from both the *Sophophora* and *Drosophila* subgenera.

Furthermore, the most basal TE clade, *TR1*, is found in 31 species, while *TR2*, one of the more derived clades, includes elements from 68 species (**Figure 2A**).

There are two evolutionary scenarios that could explain these observations: (1) the divergence of the majority of telomeric TE clades predated the common ancestor of the *Drosophila* genus and different TE clades were subsequently lost from different *Drosophila* lineages or (2) divergence of the TE clades accompanied the divergence of the genus, followed by frequent horizontal transfer of TE clades among *Drosophila* species groups. To assess the evidence for these two scenarios, we performed gene tree/species tree reconciliation (see Methods) to infer duplication (in the case of TEs, this would be divergence of an ancestral TE family into two related families within a single host lineage), transfer, and loss events for the telomeric TEs. To more easily visualize the reconciliation results, we focused on 9 major species clades (46) and pruned the species and gene trees to include the minimum number of species necessary to account for the presence of each *TR* clade within each species clade (**Figure 2B**). Surprisingly, we found that the inferred rate of horizontal transfer (HT) exceeded the inferred rates for both duplications and losses (HT: 24, duplications: 6, losses: 9). We repeated this analysis on the unpruned species and gene trees, which infers a total of 81 HT events, 16 duplication events, and 91 losses. For comparison, we ran the same analysis pipeline on a set of 465 non-telomeric Jockey TEs recently identified in *Drosophila* (48). The rate of horizontal transfer for telomeric TEs is approximately 40% higher than that of non-telomeric TEs, a difference that is highly significant (Wilcoxon test P < 2.2e-16, Figure S8).

In summary, our reconciliation analysis suggests that the six major telomeric TE clades originated within different *Drosophila* lineages at different timepoints during the diversification of the genus and subsequently expanded their host range via horizontal transfer. The predominance of both horizontal transfer and lineage-specific extinction during the evolutionary diversification of telomeric TEs is consistent with a parasitic relationship between these elements and their hosts. This relationship is similar to what has been described for facultative bacterial endosymbionts like *Wolbachia*, *Rickettsiella*, and *Spiroplasma*, all of which have been shown to undergo frequent horizontal transfer among host species while also selfishly enhancing their own fitness at the expense of their host through strategies such as cytoplasmic incompatibility and male killing (49–52). These facultative endosymbionts are distinct from obligate mutualists, like *Buchnera*, which are vertically transmitted and provide essential nutrients to their aphid hosts, with whom they have codiverged for over 200 million years (43, 53).

### Convergent evolution of telomere-specialization

Telomeres can act as a “safe harbor” where TEs can insert without deleterious consequences. Indeed, transposon insertions have been identified within the canonical telomere repeats in species of fungi, silk moths, and plants (54–57). One might therefore expect other transposons to target their insertions to *Drosophila* telomeres, yet our ORF1 and ORF2 trees (**Figure S3**) suggest a single evolutionary origin for all *Drosophila* telomeric TEs, consistent with prior work (11, 24). To further assess this possibility, we asked whether there were *Jockey* family TEs from the non-telomeric clade that form head to tail arrays at the telomeric regions of any species included in our analysis. Excitingly, we identified *Jockey* retrotransposons from the non-telomeric clade at the telomeres of six species: *D. cardini*, *D. funebris*, *D. littoralis*, *D. virilis*, *Scaptomyza hsui*, and *Scaptomyza pallida* (**Figure 3**). Interestingly, these TEs form a monophyletic clade within the larger clade of non-telomeric *Jockey* elements, despite their appearance in species that are only distantly related. For example, *D. cardini*, *D. funebris*, and *S. hsui/pallida* diverged from each other more than 20 mya (58). The presence of these closely related TEs in distantly related species is indicative of horizontal transfer (**Figure 3, Figure S9**). Hereafter, we refer to this as the NTT clade (Non-telomeric clade TEs at Telomeres).

**Figure 3.**
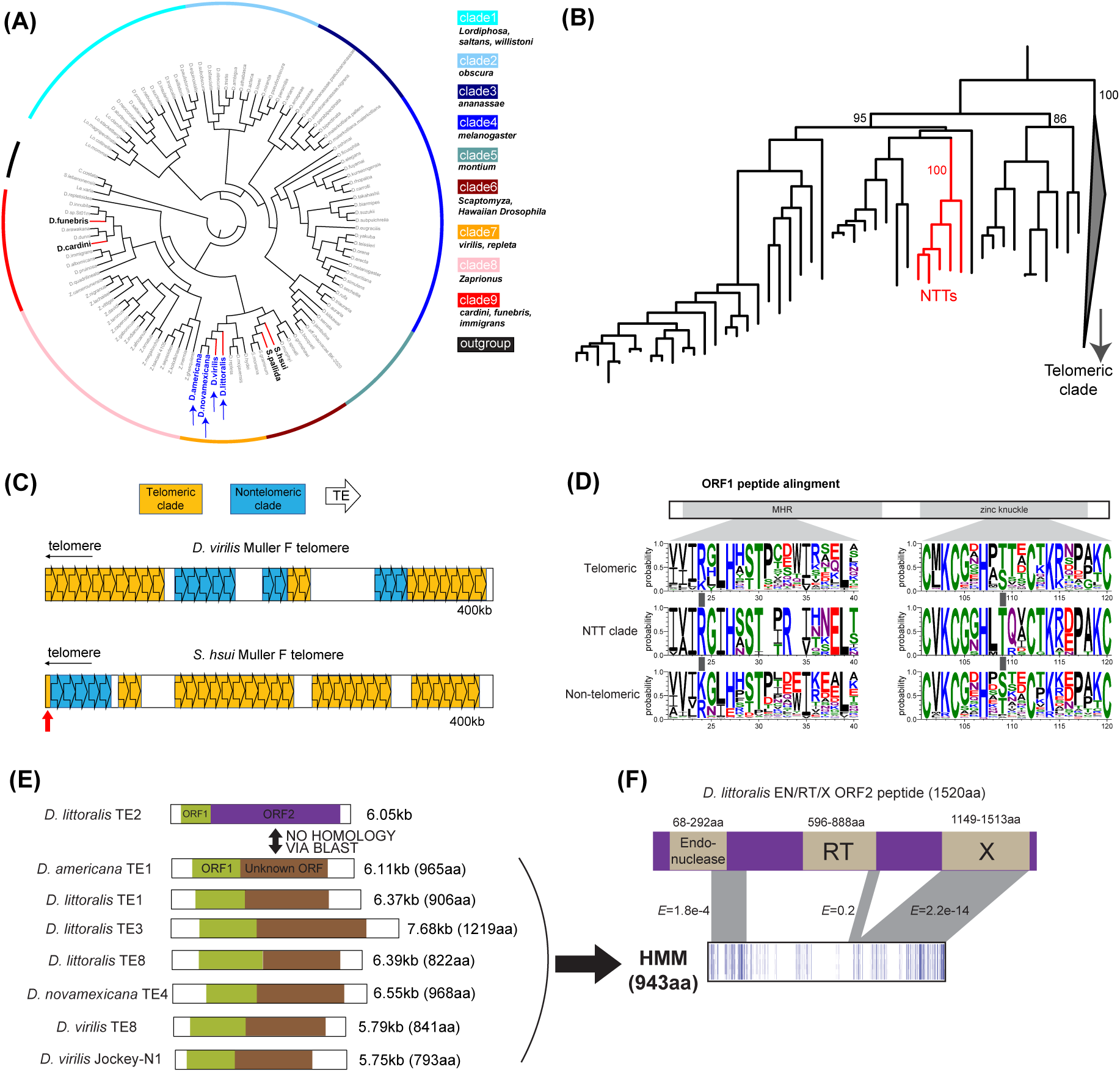
Convergence and neofunctionalization in telomeric TE families. (A) Species tree showing the six species harboring non-telomeric clade TEs at their telomeres (NTTs, red branches) and the four species carrying potentially neo-functionalized ORF2 (blue arrows). The outer circle colors delineate major species clades. (B) Pruned TE tree showing that the NTT families (red branches) form a monophyletic subclade within the larger clade of non-telomeric Jockey elements, inferred from ORF2 peptide sequences (C) Schematic examples of two species (*D. virilis* and *S. hsui*) showing telomeric regions composed of head-to-tail arrays of telomeric TEs and NTTs. In *S. hsui*, we found a fragmented TE sequence (red arrow) at the telomeric end of the contig whose partial ORF2 sequence placed it within the telomeric clade. (D) Sequence logos created from an ORF1 peptide multiple sequence alignment, focusing on the MHR and zinc knuckle motifs within the ORF1 peptide. Dark gray rectangles mark the two residues (one in each motif) that are common among both NTT and telomeric elements, but less common among other non-telomeric elements. At the first position, arginine is found at 76% of telomeric ORF1 sequences, 100% of NTT ORF1 sequences, and only 39% of non-telomeric ORF1 sequences. At the second position, threonine is found at 53% of telomeric ORF1 sequences, 100% of NTT ORF1 sequences, and only 31% of non-telomeric ORF1 sequences. (E) Seven telomeric TEs from members of the *virilis* species group (see Figure 3A) contain an ORF2 (brown box) that shows no significant homology to the ORF2 (purple box) of other telomeric TE families from this same group via peptide BLAST. Lengths correspond to the full length TE in basepairs (bp) and the ORF2-encoded peptide in amino acids (aa). (F) A profile HMM created from the seven unusual ORF2 sequences shows homology to several regions of the ORF2 from a *D. littoralis* telomeric TE that also contains the canonical endonuclease and RT domains usually found in Jockey clade ORF2 peptides, plus a domain of unknown function termed X. The unusual ORF2 sequences in Panel E may represent a case of neofunctionalization, where the ancestral function provided by the endonuclease and RT domains has been lost and replaced by a currently unknown function.

These elements are not found elsewhere in the genomes of any of the six species listed above, consistent with telomere specialization; however, in all six species, their arrays are found in between and/or adjacent to arrays of telomeric-clade *Jockey* TEs (**Figure 3C**). Their genomic location suggests that these elements have acquired the ability to target their insertions to the telomeric regions of the genome. Nevertheless, it remains unclear whether they are able to act as *bona fide* telomeres (i.e. by mobilizing directly to chromosome ends) or if they are instead parasitizing the telomeric niche by homing in on pre-existing telomeric TE arrays.

We next sought to identify amino acid substitutions in the NTT elements that might be associated with the ability to target their insertions to chromosome ends. In *D. melanogaster*, the ORF1 protein from the telomeric *HeT-A* TE has been shown to physically interact with the ORF1 proteins from the other two telomeric TE families, *TART* and *TAHRE*, directing their entry into the nucleus and their localization at the telomeres (15, 59). The region of the *HeT-A* ORF1 peptide that spans the MHR and zinc knuckle motifs was found to be required for this collaborative targeting process (15, 59). We compared this region of ORF1 from elements in the NTT clade to elements from the telomeric clade as well as the non-telomeric clade of *Drosophila Jockey* retrotransposons. Within this region of ∼137 amino acids, we identified 2 locations (positions 24 and 109) where the most common amino acid in the NTT clade was the same as that of the telomeric clade, but differed from the most common amino acid in other non-telomeric elements, to which the NTT clade is more closely related (**Figure 3D**). The number of sites showing such a pattern is significantly more than expected by chance (permutation test *P* = 0.003, see Methods). This pattern of amino acid evolution raises the possibility that the ORF1 proteins encoded by members of the NTT clade have evolved to allow targeting of their insertions to telomeres either directly or indirectly via interaction with the ORF1 proteins of telomeric elements.

### Potential neofunctionalization of ORF2

The ORF2 of all LINE elements, including mammalian LINE-1, contains, at a minimum, an endonuclease and a reverse transcriptase (RT) domain (60). The domain organization of ORF2 within the *Jockey* clade of LINEs, which includes all *Drosophila* telomeric TEs, is similar to that of LINE-1 elements, with the endonuclease domain preceding the RT domain (60, 61). The presence of a conserved endonuclease domain within the ORF2 carried by *Drosophila* telomeric TEs is surprising considering that transposition to chromosome ends should not require DNA nicking (16). In order to better characterize the extent of ORF2 domain conservation across *Drosophila* telomeric TEs, we used InterPro to identify conserved domains in the ORF2 sequences from our library of telomeric TEs. Surprisingly, we identified several telomeric TE families from *virilis* group species whose ORF2 peptides lack both the endonuclease and RT domains. According to InterPro, these unusual ORFs contain no conserved domains at all, aside from a single disordered region. The first ORF in these TEs is clearly homologous to the ORF1 protein found in other *Drosophilia* telomeric TEs, which is how these elements were classified as telomeric TEs in the first place. However, a peptide BLAST search was unable to detect any significant amino acid similarity between these unusual ORF2 peptides and the canonical ORF2 peptides from other telomeric TEs (**Figure 3E**).

In total, we identified six telomeric TE families carrying this unusual ORF from the following species: *D. virilis*, *D. americana*, *D. novamexicana*, and *D. littoralis*. The peptides encoded by these ORFs range in size from 793 to 1219 amino acids and they range from 30% to 96% identical among the seven TE families (**Figure 3E, Figure S10**). We were unable to identify homologs of these peptides outside of the above species, despite employing sensitive HMM-based search strategies (see Methods). We calculated the *dN*/*dS* ratio for all pairs of sequences and found that, in 20 out of 21 pairwise comparisons, there is statistically significant evidence of purifying selection, which suggests that the ORFs encode functional proteins (**Table S4**). The pair that did not reach significance are highly similar to one another resulting in a lack of power to detect purifying selection (**Table S4**).

In addition to the conserved endonuclease and RT domains, the ORF2 of some *Drosophila* telomeric TEs carry an additional ∼400 amino acid, glutamine rich region at the C terminus, referred to as the “X domain” (62). We hypothesized that perhaps the unusual ORF that we discovered was derived from this X domain, having lost both the endonuclease and RT domains. To test this hypothesis, we used a Hidden Markov Model (HMM) approach (see Methods) to compare the 7 unusual ORF2 peptides identified here to all telomeric TE ORF2 peptides containing the endonuclease, RT, and X domains (hereafter referred to as EN/RT/X ORF2 peptides). These searches revealed several regions of homology between a *D. littoralis* EN/RT/X ORF2 peptide and our HMM (**Figure 3F**). Small portions of both the endonuclease and RT domains in the *D. littoralis* EN/RT/X peptide show weak homology with our HMM (HMMER E-value = 1.8e-4 and 0.2, respectively). Interestingly, there is much stronger homology between the C terminus of the HMM and the X domain of this peptide (HMMER E-value = 2.2e-14).

These results suggest that the unusual ORF2 peptides we discovered are in fact derived from the canonical ORF2. However, the endonuclease and RT domain regions have evolved so rapidly that they are no longer recognizable as conserved domains and may no longer retain their ancestral function, raising the possibility that this ORF has acquired a novel function in these species. This represents, to our knowledge, the only known example of a LINE ORF2 peptide that lacks both RT and EN domains.

### Gene capture by telomeric transposons

In plants, gene capture may allow TEs to evade host silencing by forcing the host to reduce the efficiency of TE suppression mechanisms in order to avoid self-silencing (63, 64). Similarly, our prior work showed that the *D. melanogaster* telomeric *TART* element has captured a fragment of the host piRNA pathway gene *nxf2* and that *TART*-derived piRNAs likely target *nxf2* for suppression (22). This anti-silencing strategy imposes a cost to the host genome while benefitting the TE, consistent with *TART* acting as a parasite rather than a mutualist. We therefore sought to determine if gene capture occurred in the telomeric transposons of other *Drosophila* species by searching the full length consensus of all *Drosophila* telomeric TEs identified here for homology to *D. melanogaster* peptides. After manual curation of all BLAST hits (see Methods, **Figure S11**), we identified 20 telomeric TE families that show evidence of gene capture (**Table S5**), which represents at least 9 independent gene capture events across the genus, including the previously described capture of *nxf2* (**Figure 4A**). Surprisingly, eight of the nine events involve capture of known piRNA pathway genes: the capture of *nxf2* by *D. melanogaster* plus seven additional capture events involving either *piwi* or *aubergine* (*aub*), both of which are members of the Piwi subfamily of Argonaute proteins that bind piRNAs and are required for transposon silencing (32, 65–67).

**Figure 4.**
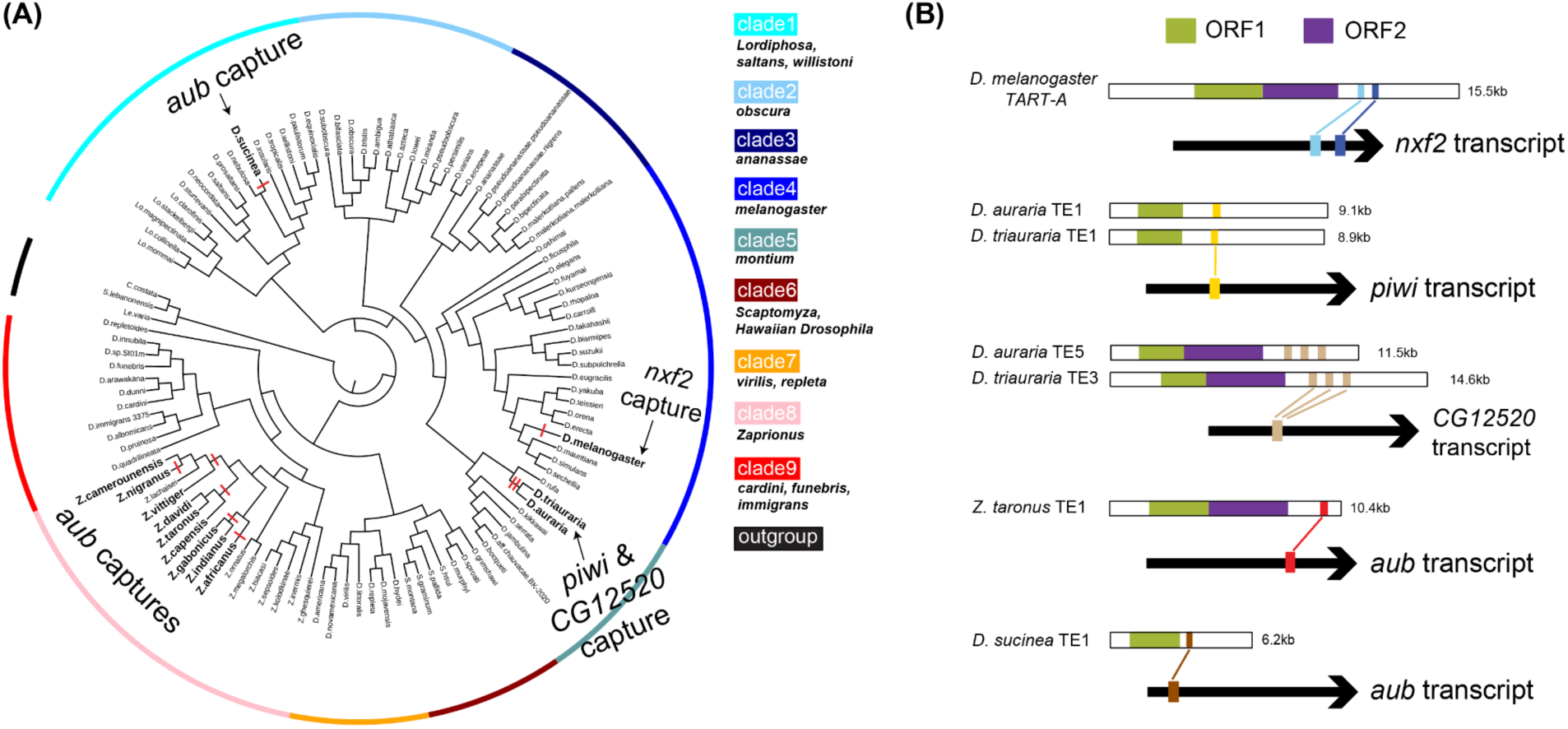
**Repeated capture of piRNA pathway genes by telomeric TEs**. (A) Species tree showing independent gene capture events (red thick lines) at the nodes. Labels indicate the gene that was captured and the outer circle is colored by major species clade. (B) Diagrams showing the location of the captured sequence within the full length TE consensus sequence and within the host gene. Green and purple boxes indicate the locations of ORF1 and ORF2, respectively. TEs lacking a purple box have lost ORF2. Note that the 5’ UTR of *TART-A* is copied from a portion of the 3’ UTR during replication, thus the 3’UTR of *TART-A* also carries *nxf2* sequences (22), but they are not shown here for clarity.

We find that a fragment of *piwi* was captured by a telomeric TE family present in both *D. auraria* and *D. triauraria* and fragments of *aub* were captured by a telomeric TE family in *D. sucinea* and nine different species within the *vittiger* subgroup of the *Zaprionus* clade. The other novel capture event involves another telomeric TE family from *D. auraria* and *D. triauraria,* which carries fragments from a homolog of the F-box domain containing *D. melanogaster* gene *CG12520*. F-box proteins are known to provide substrate specificity to the SCF ubiquitin ligase complex (68). *CG12520* is expressed in the female germline (69) and knockdown of *CG12520* causes a very modest upregulation of two telomeric TEs (70). This gene is largely uncharacterized; for example, its binding substrate remains unknown (71). Across all gene capture TE families, the length of the captured gene fragments ranges from 140bp to 235bp and their similarity to their host gene ranges from a percent identity of 76.4% to 94.82% (**Figure 4B**, **Table S5**).

Five of the nine independent gene capture events occurred in the *Zaprionus* clade. Out of 17 *Zaprionus* species with sequenced genomes, we find ten species harboring one or more telomeric TEs that contain *aub* gene fragments (**Figure 5A**). In total, four different regions of the *aub* gene were captured by different TE lineages within this group (**Figure 5A**). Some of these telomeric TEs, such as those from *Z. africanus*, *Z. gabonicus*, and *Z. indianus*, captured two different regions of the *aub* gene (**Figure 5A**). The captured *aub* fragments were subsequently amplified within these TEs, up to 7 copies in the *Z. africanus* TE and 8 copies in *Z. gabonicus* and *Z. indianus* TEs (**Figure 5A**).

**Figure 5.**
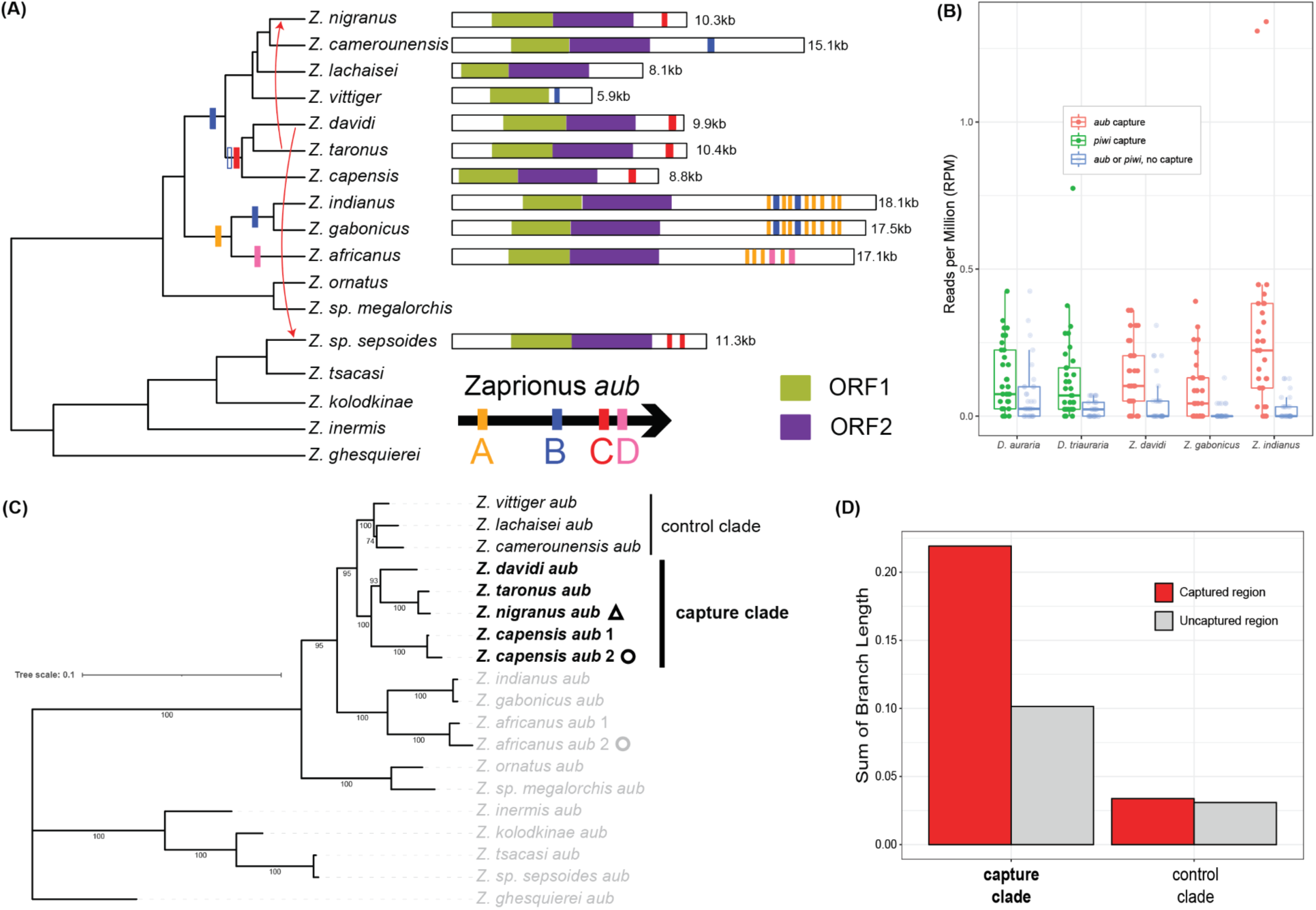
**Multiple captures of *aubergine* by telomeric TEs in *Zaprionus*. (**A) *Zaprionus* species tree showing independent gene capture events. The rectangles are colored based on the *aub* region that was captured. Diagrams show the location of the captured region within the full length TE. (B) Normalized piRNA abundance from the captured host gene (red = *aubergine*, green = *piwi*) calculated in 80 bp sliding windows across the gene transcript. *Aub* is used as a control (blue) in species where *piwi* was captured and *piwi* is used as a control in species where *aub* was captured. The control gene produces significantly fewer piRNAs in all within-species comparisons (Wilcoxon test *P* < 0.05 in all cases, see individual p-values below). (C) Gene tree inferred using *aub* coding sequence. The clades analyzed in panel D are indicated by vertical bars. (D) Comparison of rate of evolution of the captured (red) portion of *aubergine* versus the uncaptured portion. In the species where telomeric TEs have captured part of *aub* (capture clade), the homologous sequence in the *aub* gene is evolving approximately twice as fast as the uncaptured portion of the gene (Fisher’s Exact Test *P* = 2.7e-5). In species where this region has not been captured, it evolves at a similar rate as the uncaptured portion of the gene (Fisher’s Exact Test *P* = 0.64). P-values for Panel B Wilcoxon Tests: *D. auraria P* = 0.026, *D. triauraria P* = 1.3e-4, *Z. davidi P* = 3.0e-4, *Z. gabonicus P* = 6.3e-4, *Z. indianus P* = 5.2e-8.

It is possible that these seemingly independent gene capture events actually originated via a single capture event that included a large portion of the *aub* gene, followed by independent deletions of *aub* sequence in different TE lineages. Indeed, the *Zaprionus* TE families that carry *aub* sequence form a monophyletic clade in the ORF2 TE tree (**Figure S12**). We tested this prediction by reconstructing the evolutionary history of each capture event (**Figure S12**). We created gene trees using alignments containing the captured *aub*-like TE sequences along with their homologous sequences from the *aub* gene of each *Zaprionus* species. These trees confirmed our initial inference that five independent capture events occurred within *Zaprionus* and allowed us to assign each event to a node in the *Zaprionus* species tree (**Figure 5A**). Unexpectedly, we find that region B of *aub* was captured in the ancestor of *Z. vittiger* and *Z. capensis* and subsequently lost in the *Z. davidi* / *Z.taronus* / *Z.capensis* clade, whose TEs now carry region C of *aub* (**Figure 5A, Figure S12**). We also note that region B of *aub* was captured independently by two different *Zaprionus* clades; however, the captured regions overlap but do not share the same boundaries.

The repeated capture of piRNA pathway genes by *Drosophila* telomeric TEs suggests that acquisition of these gene fragments may increase TE fitness. We previously found evidence that antisense piRNAs produced from the *nxf2*-like region of *TART* are capable of targeting the *nxf2* host gene for suppression, consistent with an anti-silencing strategy to escape host suppression (22). To determine if a similar scenario is occurring in these other species, we performed small RNA sequencing from ovaries for five species where telomeric TEs had captured either *aub* or *piwi* as well as 4 related species where gene capture had not occurred to use as controls. The median identity between captured sequence and host gene is ∼90% across all gene capture TEs (**Table S5**). This identity is noticeably smaller for two gene capture cases: *CG12520* in *D. auraria*/*triauraria* (∼79% identity) and *aub* in *Z. sp. sepsoides* (∼82% identity)(**Table S5**). We find almost a complete lack of piRNAs mapping to the host genes in these latter two cases of elevated divergence (**Figure S13**), suggesting that the similarity between captured sequence and host gene is too low to direct host gene targeting by TE-derived piRNAs.

In the species with high sequence identity between the captured sequence and the host gene, we found small RNAs from both the gene capture telomeric TEs (**Figure S14**) and the *aub* (or *piwi*) host genes whose lengths and 5’ U bias are consistent with piRNAs (**Figure S15**). Furthermore, in species where *aub* (but not *piwi*) was captured, the abundance of piRNAs from *aub* is significantly larger than from *piwi* while the reverse is true for species where *piwi* (but not *aub*) was captured (**Figure 5B**). Overall, piRNA abundance from the captured host gene is significantly larger in the species harboring gene capture TEs compared to species whose TEs have not captured any host gene (Wilcoxon test P = 0.0036, **Figure S16**), consistent with the host gene being targeted by piRNAs derived from the gene capture region of the telomeric TE. Across all species harboring gene capture TEs, the telomeric TE families that have captured a host gene fragment have significantly higher copy numbers compared to the other non-capture telomeric TEs in the same genome (paired Wilcoxon test, one-sided *P* = 0.02, Figure S17), suggesting that gene capture increases TE fitness.

If *aub* is being targeted for suppression by TE-derived piRNAs, we expect to see accelerated evolution in the corresponding region of *aub*, which would reduce piRNA targeting by decreasing the similarity between *aub* and the *aub*-like sequences within the telomeric TE. We focused on the clade of *aub* with sequences from the following species: *Z. nigranus*, *taronus*, *davidi*, and *capensis*, since the telomeric TEs in all of these species have captured the same single region of *aub*, and the captured sequences have not been amplified within the TE (**Figure 5A**). We find that the region of *aub* that was captured by TEs is evolving approximately twice as fast as the uncaptured portion of the gene, based on a comparison of trees constructed from the captured region of *aub* versus *aub* sequence that was not captured by any *Zaprionus* TEs (Fisher’s Exact Test *P* = 2.7e-5, see Methods). As a control, we repeated this analysis on the *aub* clade from *Z. camerounensis*, *lachaisei*, and *vittiger*, whose TEs have not captured this region of *aub*. In this case, there was no difference in the rate of evolution (Fisher’s Exact Test *P* = 0.64, **Figure 5D**).

We also find that *aub* has undergone two independent duplication events in *Zaprionus*, once in the *Z. capensis* lineage and once in *Z. africanus* (**Figure 5C**). This finding alone is remarkable given that a previous survey of 39 Dipteran species (including 22 species of *Drosophila*) identified only a single duplication event involving *aub* since its origin over 150 million years ago, which occurred in stalk-eyed flies (72). Notably, both *Zaprionus aub* duplications occurred in species with telomeric TEs that have captured an *aub* fragment (**Figure 5A**), raising the possibility that the duplications were selected for as a means to increase *aub* dosage in reponse to it being targeted by TE-derived piRNAs.

In addition to duplications, we also find strong support for introgression of *aub* from *Z. taronus* to *Z. nigranus*, likely due to recent gene flow between these two species (**Figure 5C, Figure S18, Figure S19**). This transfer of *aub* from *taronus* to *nigranus* potentially set the stage for a subsequent invasion of the *nigranus* genome by *taronus* telomeric TEs, whose *aub*-like sequences are 92.26% identical to those of the *taronus aub* gene. Indeed, two telomeric TEs, both of which carry an *aub* gene fragment, show evidence of horizontal transfer from *Z. taronus* to *Z. nigranus* (**Figure S18)**. Based on sequence similarity, we infer that the two telomeric TEs initially diverged in *Z. taronus*, with one family subsequently losing the ORF2 gene (**Figure S18)**. Synonymous divergence between *taronus* and *nigranus* copies of the *aub* gene (*Ks*=0.034) is larger than that of the ORF1 genes for the two TE families (TE-1 *Ks*=0, TE-2 *Ks*=0.023), suggesting that the transfer of *aub* from *taronus* to *nigranus* occurred first, followed by the transfer of TE-2 and, most recently, the transfer of TE-1 (**Figure S18**). Notably, these recently transferred TEs (which captured region C of *aub*, Figure 5A) have replaced the ancestral telomeric TEs in *Z. nigranus*, which carried region B of *aub*.

Thus, different lineages of gene capture TEs may compete with one another to maintain their residence in a given host genome.

Together, these results suggest that capture of piRNA pathway genes by telomeric TEs has occurred repeatedly across the *Drosophila* genus, likely as a form of counter-defense, where TEs selfishly target host suppression systems for silencing to increase their own fitness. The accelerated evolution of the region of the host gene that was captured, which was previously described for *nxf2* (22) and shown here for *aub*, suggests there is selection to avoid this targeting by “erasing” the homology between the captured sequence carried by the TE and the host gene.

## Discussion

Recent work in both *Arabidopsis* and Zebrafish has introduced the concept of transposon “addiction” where an active TE family serves an essential function for its host genome despite causing mutational damage via insertion of new TE copies (6, 7). The telomeric TEs of *Drosophila* fit nicely into this framework: they play a critical role in protecting and extending chromosome ends and, importantly, the benefit they provide to their host *depends* upon their transposition activity. However, the host genome also constrains the activity of these elements, both to ensure that transposition occurs only on chromosome ends (27) and to control telomere length as ultralong telomeres have been shown to reduce fertility (37). The TE addiction model predicts that telomeric TEs should eventually be fully co-opted by their host genome; however, our results suggest that the tension between selfish mobilization and host control has led to a prolonged period of evolution between these TEs and their host genome that is dominated both by instability and genetic conflict.

Rather than the co-speciation that one might expect from a true genetic mutualist, the telomeric TE phylogeny is dominated by horizontal transfer and lineage-specific extinction events. In fact, we find that the relationship between telomeric TEs and their host has completely unraveled at least 10 separate times across the genus, with a total of 13 *Drosophila* species (and two outgroup species) having lost telomeric TEs entirely, and thus presumably relying on an alternative, recombination-based mechanism of chromosome elongation (*i.e.* Alternative Lengthening of Telomeres (ALT) (73)). This instability may be due in part to the dynamic nature of the telomeric niche (41); however, we also find evidence of competition among telomeric TE clades. Both the TR1 and TR6 clades predate the origin of the TR2 clade, but TR2 elements are found in more than half of all species studied here, which is ∼2-fold as many species compared to TR1 and ∼10-fold the number of species carrying TR6 clade TEs. TR2 has therefore spread via horizontal transfer into lineages whose ancestral telomeres were occupied by TR1 and/or TR6 clade TEs but have since been replaced by TR2 elements.

We also find evidence of potential competition among TEs from non-telomeric *Jockey* clades for the telomeric niche. The non-telomeric clade TEs at telomeres (NTTs) we identify here are well supported as members of the non-telomeric *Jockey* clade, yet they appear to be present in head-to-tail arrays exclusively at the telomeres of six *Drosophila* species, always either adjacent to, or in between, arrays of telomeric *Jockey* clade TEs. This localization pattern suggests that these elements are able to target their insertions to the telomeric regions of chromosomes. Future work will determine whether they are able to directly mobilize to chromosome ends or if they instead only insert into pre-existing arrays of telomeric clade TEs.

In contrast to what would be expected from a genetic mutualist, we find evidence of extremely rapid evolution and potential functional innovation within a subset of telomeric TE families from several *virilis* group species. The ORF2 amino acid sequence of these TE families has evolved so rapidly that it is almost unrecognizable as an ORF2 homolog, which is especially notable given that ORF2 (as opposed to ORF1) is preferentially used for phylogenetic classification of LINEs due to its slower rate of evolution. The endonuclease and reverse transcriptase conserved domains are hallmarks of ORF2 encoded peptides in LINE non-LTR retrotransposons from humans to nematodes (60). Neither of these domains are detectable by InterPro (74) in the amino acid sequences of these unusual ORF2s. It is possible that the loss of these conserved domains is caused by degeneration or pseudogenization. However, our *dN*/*dS* analysis shows that these unusual ORF2 sequences are evolving under purifying selection. Thus, these peptides may have evolved a novel function that is different from the host DNA cleavage and reverse transcription functions provided by canonical ORF2s, which is now being maintained by purifying selection. How could these TEs continue to mobilize after loss of these two critical ORF2 functions? It is notable that the entire ORF2 is dispensable from a given TE as long as it is encoded by other related TE families, as evidenced by the repeated loss of ORF2 from many telomeric TEs. Such dispensability may have provided this unusual ORF2 the flexibility to evolve a novel function similar to the phenomenon of neofunctionalization after gene duplication (75).

Frequent horizontal transfer, competition among telomeric TE lineages, and functional innovation within certain lineages are all consistent with antagonistic evolution between telomeric TEs and their hosts. The possibility of anatagonistic coevolution is further supported by our evidence that that telomeric TEs have repeatedly captured fragments of specifically piRNA pathway genes, including *aubergine* and *piwi*, in addition to the previously described capture of *nxf2* by the *D. melanogaster* telomeric transposon *TART* (22). PiRNA abundance from a subset of these species is consistent with our previous work, suggesting that piRNAs derived from the captured gene fragment(s) embedded within the telomeric TEs are capable of targeting the host gene for suppression (22). This phenomenon thus likely represents a counter-defense or anti-silencing strategy deployed by these TEs (5, 76–80). It may seem self-destructive for a single TE family to impair the piRNA pathway, thus potentially leading to the upregulation of the majority of active TEs in the genome. However, the telomeric transposons of *D. melanogaster* are exquisitely sensitive to disruption of the piRNA pathway (70). Assuming telomeric TEs in other species are similarly sensitive, they may be able to suppress piRNA activity enough to benefit themselves without leading to upregulation of other TE families.

Similar to what we observed for *nxf2* in *D. melanogaster*, we find accelerated evolution of the *aub* gene in *Zaprionus*, specifically in species harboring *aub*-capture TEs and specifically in the region of *aub* that was captured. This process is consistent with antagonistic coevolution: the telomeric TE selfishly captures a fragment of the host gene, which decreases expression of the host gene via piRNA-mediated silencing. Host gene mutations that disrupt piRNA targeting would then be favored by natural selection, leading to accelerated evolution of the host gene in the region that was captured. The TE may respond by capturing a new region of the same gene, as we see in *Zaprionus*. Amplification of the captured gene fragment (as seen in *Z. indianus*, *Z. africanus*, and *Z. gabonicus*) may also be beneficial by allowing the TE to more efficiently acquire mutations that increase or preserve the similarity between the captured fragment and host gene via non-allelic gene conversion among the duplicated fragments (81, 82).

The well-ordered arrays of telomeric TEs found at *Drosophila* chromosome ends are the result of a complex interplay between TE families that both cooperate and compete with each other as well as the host piRNA (28–31), DNA repair (83, 84), chromosome end-capping (85), and heterochromatin maintenance (26, 86) pathways, each of which is necessary for proper telomere function. Telomeric TE lineages are likely to have co-evolved to at least some degree with the host proteins involved in these pathways to ensure the correct formation and maintenance of telomeric arrays. It is therefore remarkable that both telomeric TEs and host proteins from all three of these pathways are known to evolve rapidly (23, 87, 88). Previous work has proposed various types of genetic conflict that could explain this observation (23). Our work here adds two new possibilities: (1) the frequent horizontal transfer of telomeric TEs among species, as well as the invasion of the telomeric niche by NTTs, could require rapid host protein evolution to maintain coordination between TE mobility and telomere maintenance and (2) repeated gene capture by telomeric TE families could antagonize the piRNA pathway, leading to rapid evolution of pathway members.

Another striking outcome of our study is the degree to which convergent evolution occurs among *Drosophila* species and among telomeric TE lineages. To the extent that this convergence is a result of adaptive evolution, these events provide insight into both host and TE biology. The repeated loss of telomeric TEs from various *Drosophila* species raises the possibility that these elements impose a fitness cost to their host and that it is advantageous to be rid of them when an alternative mode of chromosome elongation is available. In terms of the TEs themselves, their frequent loss of ORF2 raises the possibility that TEs lacking this ORF are more fit than their progenitors (as long as they can utilize ORF2 peptides encoded other TE families). Similarly, the repeated capture of piRNA pathway genes by telomeric TEs suggests that gene capture increases TE fitness (see above). Horizontal transfer is emerging as a key mechanism by which TEs avoid extinction (89, 90) and our observation of repeated horizontal transfer of telomeric TEs supports such findings. Finally, our finding of convergent evolution of telomere localization is consistent with the telomeric niche acting as a “safe harbor” for TE insertions and thus, TEs with telomeric insertion biases may have increased fitness relative to other TEs.

The “transposon addiction” model envisions an intermediate period of cooperation between TE and host in the continuum between selfish TE mobilization and host co-option (or TE extinction) (7). We show that this cooperation period can persist for millions of years: Telomeric TEs originated over 60 million years ago, before the origin of the *Drosophila* genus, and remain present in the vast majority of *Drosophila* species that we surveyed in our study. However, cooperation and conflict are not mutually exclusive. Throughout the 60+ million-year period of apparent cooperation, we find numerous signs of conflict both between different telomeric TE lineages and between these TEs and their host genome. It is striking that these signatures of antagonistic evolution recurrently appear across the *Drosophila* phylogeny, showcasing the major role of convergence in the evolution of these TEs. This convergence extends beyond *Drosophila*, as well. Telomerase, the holoenzyme responsible for telomere elongation across most eukaryotes, utilizes a reverse transcriptase that likely originated from an ancient non-LTR retrotransposon (91–93). Thus, the same phenomenon of TE addiction that is currently playing out in *Drosophila* may have also occurred over a billion years ago, ultimately leading to the co-option of a retrotransposon to form the reverse transcriptase component of the holoenzyme that we now know as telomerase (91).

There are echoes of convergence in other TE addiction systems as well. For example, telomeric TEs are not the only “beneficial” TEs that are able to target their insertions to specific locations in the genome. The *G2/Jockey-3* and *ATHILA* retrotransposons that likely aid in centromere formation in *Drosophila* and *Arabidopsis*, respectively, have evolved mechanisms to target their insertions to centromeric chromatin (6, 94, 95).

Similarly, the *Drosophila R2* retrotransposon, which plays a role in the maintenance of ribosomal DNA repeats, is able to target its insertions to rDNA (96). In summary, we find that what appears to be a long-standing period of cooperation between telomeric TEs and their host genome is also rife with repeated occurrences of genetic conflict and dynamic evolution. Discovery and characterization of additional TE addiction systems will help to determine the generality of the TE addiction process observed at the *Drosophila* telomere.

## Methods

### Species

*Drosophila* species used in this study were provided by Matute lab (*Z. indianus* [RCR 17.04], *Z. gabonicus* [JD’18], *Z. davidi* [JD’18], *Z. nigranus* [18 CAR07 Z03]) and ordered from the National Drosophila Species Stock Center at Cornell (*D. auraria* [strain: 14028-0471.00] and *D. triauraria* [strain: 14028-0651.00], *Z. ghesquierei* [strain: 50000-2743.00], *Z. kolodkinae* [strain: 50000-2748.00])*. Z. taronus* [strain: 50001-1020.00] and *Z. camerounensis* [strain: 50001-1010.01] were ordered from the National Drosophila Species Stock Center at Cornell, but our sequencing data from these species suggested that they were misidentified and that the two stocks are likely to be from unsequenced species related to *Z. megalorchis* and *Z. sepsoides,* respectively. We therefore refer to these stocks as *Z. sp. megalorchis* and *Z. sp. sepsoides*.

### Data acquisition

Out of the 109 long-read genome assemblies used in this study, 106 assemblies were acquired from NCBI: *D. albomicans* (NCBI BioProject PRJNA630751) (97), *D. bifasciata* (PRJNA565796) (98), *D. innubila* (PRJNA524688) (99), *D. miranda* (PRJNA474939) (100), *D. pseudoobscura* (PRJNA596268) (101), *D. serrata* (PRJNA355616) (102), *D. suzukii* (PRJNA594550) (103), *D. triauraria* (PRJNA627893) (104), *D. mauritiana*, *D. sechellia*, and *D. simulans* (PRJNA383250) (105), *D. athabasca*, *D. lowei*, and *D. subobscura* (PRJNA545704) (106), and *D.ananassae*, *D. azteca*, *D. erecta*, *D. hydei*, *D. novamexicana*, *D. orena*, *D. persimilis*, *D. virilis*, and *Scaptodrosophila lebanonensis* (PRJNA475270). The remaining 83 NCBI assemblies were obtained from (PRJNA675888) (45). Three new assemblies were generated in our lab: *D. auraria*, *Z. sp. megalorchis* and *Z. sp. sepsoides* (see below).

### Sequencing and Genome Assemblies

Using Monarch® HMW DNA Extraction Kit for Tissue, we extracted DNA from ∼20 females of *D. auraria (*NEB T3010), ∼30 males of *Z. sp. megalorchis*, and *Z. sp. sepsoides* (NEB T3060S). We used Oxford Nanopore Technologies (ONT) SQK-LSK109 library preparation kit for *D. auraria* and the ONT SQK-LSK114 library preparation kit for the two *Zaprionus* species to construct libraries following the ONT Ligation Sequencing Kit protocol. The library of *D. auraria* was sequenced on a MinION R9.4 flow cell. Each library of two *Zaprionus* species was sequenced on a MinION R10.4.1 flow cell. Raw signal data were basecalled using the ONT Guppy software package version 4.0.15 for *D. auraria* and version 6.4.8 for the two *Zaprionus* species with default parameters.

We used Flye (version 2.8.1) (107) to assemble the *D. auraria* genome, followed by polishing with Medaka (version 1.1.3) (https://github.com/nanoporetech/medaka) and Nextpolish (version 1.3.1) (108). We removed allelic contigs using Purge Haplotigs (version 1.1.2) (109). We then performed Hi-C scaffolding of the remaining contigs using the 3D de novo assembly (3D-DNA) pipeline (version 201008). Hi-C data were generated as described in (104). The two *Zaprionus* species genomes were assembled using Flye (version 2.9.2) with the option *--no-alt-contigs*, and polished using Medaka (version 1.6.0).

### Identification of telomeric retrotransposons

We used telomeric retrotransposons of the genus *Drosophila* available on the Repbase database (TART_DVIR, HeT-A_DYAK, TART-A_DMEL, TART-B_DMEL, TART-C_DMEL, TAHRE_DMEL, and HeT-A_DMEL) as the query to run *RepeatProteinMask* (REPEATMASKER version 4.1.2) (http://www.repeatmasker.org) on each species genome with the parameters *-noLowSimple -engine ncbi*. We clustered similar DNA sequences within species for ORF1 and ORF2 genes separately using CD-HIT-EST (version 4.8.1) (110) with the parameters *-d 0 -r 0 -c 0.9 -n 8 -g 1 -T 4 -M 32000*. We aligned the DNA sequences of each cluster using MUSCLE (version 3.8.31) (111) with the parameters *-maxiters 1 -diags* and then ran PILER (version 1.0) (112) to construct consensus sequences for each cluster, removing consensi less than 600 bp in length. We used the MACSE pipeline (OMM_MACSE version 10.01) (113) with parameters *--no_prefiltering --no_postfiltering*) to generate frameshift-aware amino acid translations for ORF1 and ORF2 from each consensus sequence.

We concatenated the ORF1 amino acid sequences generated by the MACSE pipeline from all species, additionally adding the following known telomeric ORF1 peptides: TART_DVIR, HeT-A_DYAK, TART-A_DMEL, TART-B_DMEL, TART-C_DMEL, TAHRE_DMEL, and HeT-A_DMEL and the following known non-telomeric ORF1 sequences: Juan_DMEL, Jockey_DMEL, Doc_DMEL, and F-element_DMEL. We did the same thing for ORF2 (minus the HeT-A peptides because this TE family lacks ORF2) and ran MAFFT (version 7.471) (114) with *--auto* to generate ORF1 and ORF2 amino acid multiple sequence alignments. We trimmed the aligned sequences using CLIPKIT (version 1.1.5) (115) with parameters *-m kpic-gappy -g 0.5* for ORF1 and *-m kpic-gappy -g 0.6* for ORF2. We removed the trimmed sequences with more than 20% and 10% gaps, for ORF1 and ORF2, respectively. We constructed ORF1 and ORF2 gene trees using IQ-TREE (version 1.6.12) (116) with parameters *-m TEST -abayes -bb 1000*.

We used an unbiased approach to define the telomeric clade for the ORF1 and ORF2 gene trees by calculating branch lengths between each novel ORF1/ORF2 gene and the known telomeric and non-telomeric ORF1/ORF2 genes included in our unrooted trees (see Github for code). The novel ORF1 and ORF2 genes that showed shorter branch lengths to their known telomeric homologs (compared to their non-telomeric homologs) were assigned as putative telomeric clade peptides. This approach resulted in a single monophyletic clade of putative telomeric ORF1 and ORF2 genes (**Figure S3**).

We next sought to identify the genomic locations of each candidate telomeric ORF1 and ORF2 gene. The telomeric clade ORF1 and ORF2 consensus DNA sequences from each species were used as a custom library when running RepeatMasker (version 4.1.2) with parameters *-no_is -norna -nolow -engine ncbi*, retaining matches with <= 3% divergence from their consensus. We used the RepeatMasker hits as starting points to generate manually curated, full-length telomeric TE consensi. We identified arrays of RepeatMasker hits at contig ends (or within short unplaced contigs) and generated a dot plot of the array sequence using YASS, a genomic similarity search web tool (117). We manually examined the dotplots to identify the boundaries of the monomers that compose the head-to-tail TE arrays. We then compared the monomers from the same TE family to identify the longest, unfragmented, representative sequence for each TE family. For species with Oxford Nanopore data, we aligned the nanopore genomic sequencing data to the representative TE sequence and performed one round of polishing using MEDAKA (version 1.6.0). All consensi (except those from species lacking Illumina data, see **Table S6**) were then further iteratively polished using PILON (version 1.24) (118) with Illumina data until no new changes were made to the consensus. We searched each consensus for intact ORF1 and ORF2 open reading frames and performed an additional round of polishing using PILON with Illumina data (where available, see **Table S6**) when the consensus sequence contained fragmented ORFs.

To estimate the genomic copy number of each telomeric TE family, we aligned the long-read genomic sequencing data from the host species to the TE conseni and divided the median TE per-base read depth by the median whole-genome read depth (averaged across 100bp windows).

### Identification of the telomeric retrotransposons with host gene capture

We used *D. melanogaster* amino acid sequences (longest isoform per gene, r6.42 annotation) as queries and the DNA consensus sequences of all species’ telomeric transposons as the database to run TBLASTN (version 2.10.1) (119) with parameters *-outfmt 6 -evalue 1e-5*. For each TBLASTN hit, we determined whether there was significant homology between the telomeric TE and its host species gene at the DNA level by running BLASTN (version 2.10.1) with parameters *-outfmt 6 -evalue 1e-3*. To search for older gene capture events we also ran BLASTN with parameters *-outfmt 6 - word_size 4* to identify host gene/TE sequences with higher levels of sequence divergence.

### Phylogenetic reconstruction (species, telomeric retrotransposons, and their reconciliation)

We used full-length ORF1 and ORF2 sequences from our curated telomeric TE consensi to create telomeric TE gene trees. We additionally added the following known non-telomeric Jockey clade ORFs from RepBase TE families to use as outgroup sequences: Jockey_DMEL, Jockey-1_DEr, Jockey-1_DEu, Jockey-1_DF, Jockey-1_DGri, Jockey-1_DK, Jockey-1_DRh, Jockey-1_DT, Jockey-1_DVi, Jockey-1_DWi, Jockey-1_DYa, Jockey-10_DAn, Jockey-10_DRh, Jockey-11_DAn, Jockey-11B_DBp, Jockey-14_DBp, Jockey-2_DEl, Jockey-2_DEu, Jockey-2_DRh, Jockey-2_DT, Jockey-3_DEu, Jockey-3_DF, Jockey-3_DK, Jockey-3_DRh, Jockey-3_DVi, Jockey-4_DEl, Jockey-4_DEu, Jockey-4_DF, Jockey-4_DPer, Jockey-4_DVi, Jockey-5_DAn, Jockey-5_DBp, Jockey-5_DTa, Jockey-6_DAn, Jockey-6_DEl, Jockey-6_DEu, Jockey-6_DF, Jockey-7_DAn, Jockey-7_DF, Jockey-8_DAn. We aligned the ORF peptides using MAFFT (version 7.471) and trimmed the alignments using CLIPKIT (version 1.1.5) with parameters *-m kpic-gappy -g 0.1*. We removed sequences with > 20% gaps from each ORF alignment and then created the telomeric ORF1 and ORF2 gene trees using IQ-TREE (version 1.6.12) with parameters *-m TEST -abayes -bb 1000* and rooted the trees using the non-telomeric outgroup sequences. All tree figures were created using the iTOL tool (120).

To reconcile the above TE ORF1 and ORF2 trees with the *Drosophila* species tree, we first pruned the dated species tree from (46), keeping only the species studied here. We then ran *Ranger-DTL-Dated* (version 2.0) (121) with default parameters to perform the gene tree/species tree reconciliation. We calculated support values for each reconciliation event using the *AggregateRanger* program on the output of 1000 *Ranger-DTL-Dated* reconciliation runs.

We generated two novel *de novo* genome assemblies for this study: *Z. sp. megalorchis* and *Z. sp. sepsoides*. We placed these species within the *Zaprionus* species tree using the same methods as described in (46). Briefly, we identified single-copy orthologs from each *Zaprionus* species genome assembly by running BUSCO (version 5.4.7) (122) with the Diptera data set of the OrthoDB v10 release (diptera_odb10) (123) using default parameters. We used IQ-TREE (version 1.6.12) to obtain the gene trees for each single-copy ortholog, as described in (46). The gene trees were used to infer the *Zaprionus* species tree using ASTRAL (version 5.7.8) (124) with default parameters.

To test whether the NTT clade shows more telomeric TE-like amino acids than expected by chance, we permuted the assignment of TEs to the NTT clade by shuffling the ids of all non-telomeric TEs 1000 times.

To make the *aub* gene tree for *Zaprionus* species, we first built *aub* gene models in each species using *genBlastG* (125) with parameters *-e 1e-5 -g T -r 3 -c 0.6* and the *D. melanogaster* Aub peptide sequence as a query. We then used MAFFT (version 7.5.20) with *–auto* to align the *aub* coding sequences and IQ-TREE (version 1.6.12) with parameters *-m TEST -bb 1000* to infer the gene trees.

To test for accelerated evolution of the *aub* gene in *Zaprionus* we first identified all regions of *aub* coding sequence that were not captured by any *Zaprionus* species (i.e. uncaptured sequence) and concatenated these regions together for each *aub* gene in each species. We aligned these sequences using MAFFT (version 7.520) with *–auto*. We then extracted two sequence subsets from this alignment: those from *Z. nigranus*, *taronus*, *davidi*, and *capensis* (capture clade), and those from *Z. camerounensis*, *lachaisei*, and *vittiger* (control clade). We added *Z. inermis aub* sequence to each sequence subset as an outgroup and used ModelFinder from IQ-TREE which identified “TPM2+G4” as the best-fitting DNA substitution model for both sequence subsets. We then inferred gene trees for both the capture and control sequence subsets from the uncaptured *aub* sequence using IQ-TREE, specifying *Z. inermis* as the outgroup and “TPM2+G4” as the substitution model and summed the branch lengths of the resulting trees.

We then repeated the above procedure using Region C of *aub*, which was captured by the *Z. nigranus*, *taronus*, *davidi*, and *capensis* telomeric TEs, providing the corresponding uncaptured treefile as a constraint tree topology. We then compared the summed branch lengths between the captured and uncaptured *aub* regions, for both the capture clade and the control clade. We used Fisher’s Exact test to compare the proportion of substitutions among all sites between captured and uncaptured sequence subsets.

To test for introgression between *Z. taronus* and *Z. nigranus*, we downloaded gene trees from BUSCO single-copy orthologs generated in a previous study (46). Across all ∼2500 trees, we counted the number of times we observed the following species pairs as sister taxa, which all share the same most recent common ancestor, according to the species tree in (46): *Z. taronus* vs *Z. nigranus* (n=71), *Z. davidi* vs *Z. nigranus* (n=9), *Z. capensis* vs *Z. nigranus* (n=12), *Z. taronus* vs *Z. vittiger* (n=11), *Z. taronus* vs *Z. camerounensis* (n=6), *Z. taronus* vs *Z. lachaisei* (n=8). Overall, 2.77% of gene trees show *Z. taronus* and *Z. nigranus* as sister taxa compared to an average of 0.35% support for the other pairs shown above.

### Small RNA sequencing

For 8 *Zaprionus* species plus *D. auraria* and *D. triauraria*, we extracted small RNAs from 10-24 pairs of ovaries using the TraPR Small RNA Isolation and Library Prep Kit (Lexogen Inc, 135.08) (126), following the kit protocol. The small RNA libraries were sent to Novogene Corporation Inc. for NovaSeq SE50 sequencing.

We used Trim Galore (version 0.6.10) (https://github.com/FelixKrueger/TrimGalore) to trim adapter sequences from raw small RNA reads and SortMeRNA (version 4.3.6) (127) to remove reads arising from tRNAs and rRNAs. TRNA genes were predicted by tRNAscan-SE (version 2.0.11) (128) and rRNA databases were generated by blastn searches with *D. melanogaster* rRNA queries. The filtered small RNA reads were aligned using ShortStack (version 3.8.5) (129) with the parameters *--dicermin 20 -- dicermax 35 --mismatches 2*.

### Detection of piRNA reads mapped to telomeric TEs and host genes, *piwi* and *aub*

We sought to quantify piRNA abundance from each telomeric TE family, as well as the host genes that were captured by telomeric TEs, excluding the captured region of the host gene to avoid cross-mapping of TE-derived piRNAs to the host gene. To do this, we masked the telomeric TEs and host genes (i.e. *piwi*, *aub*, and *CG12520*) in the appropriate species genome using *RepeatMasker* (REPEATMASKER version 4.1.2) with parameters *-no_is -norna -nolow -pa 4 -engine ncbi*. We then created a custom reference genome by appending consensus sequences of each telomeric TE family as well as mRNA sequences of the longest isoform of the captured host gene(s) (where the captured region was masked using BEDTOOLS, version 2.25.0 (130)), to the masked genome assembly. Next, we used ShortStack (version 3.8.5) with the parameters *--dicermin 20 --dicermax 35 --mismatches 2* to align the small RNA reads to the custom reference genome. We retained reads with lengths between 23 bp and 30 bp and used BEDTOOLS (version 2.25.0) to calculate coverage of sense and anti-sense alignments on the telomeric TEs and the relevant host transcript(s) (i.e. *piwi*, *aub*, and/or *CG12520*).

### dn/ds (Ka/Ks) analysis

We used MEGA X (131) to align coding sequences and used *KaKs_Calculator 2.0* (132) to calculate the ratio of the number of nonsynonymous substitutions per nonsynonymous site to the number of synonymous substitutions per synonymous site.

Fisher’s Exact Test was used (within *KaKs_Calculator*) to test the null hypothesis of neutral evolution (i.e. equal rates of synonymous versus non-synonymous substitutions).

### HMM searches

We initially used an alignment of all seven unusual ORF2 sequences as a query to perform sensitive HMM-HMM sequence searches against the *UniRef30_2023_02* database on the MPI Bioinformatics Toolkit webserver (133) using HHblits (134) but found no significant hits outside of our query species. We then aligned the seven unusual ORF2 amino acid sequences using MAFFT (version 7.520) and created an HMM profile from the alignment using hmmbuild from HMMER3 (version 3.4)(135). We compiled a database of all ORF2 peptides in *D. virilis*, *D. americana*, *D. novamexicana*, and *D. littoralis* telomeric TEs that contained the endonuclease, RT, and X domains and searched this database with our HMM profile as the query using hmmsearch from HMMER3.

## Data availability

All code used for this study, as well as the manually curated telomeric TE consensus sequences, are available on Github: https://github.com/jaehakson/DrosophilaTelomericRetrotransposons. The data generated for this project include whole genome sequencing data and genome assemblies for *D. auraria*, Z. *sp.* megalorchis, and Z. *sp.* sepsoides as well as small RNA sequencing data from 7 Zaprionus species (*Z. davidi, Z. gabonicus, Z. ghesquierei, Z. indianus, Z. kolodkinae, Z. sp. megalorchis, and Z. sp. sepsoides*), *D. auraria* and *D. triauraria.* All data are available under the NCBI BioProject PRJNA1183390.

## Acknowledgements

The authors acknowledge the Office of Advanced Research Computing (OARC) at Rutgers, The State University of New Jersey for providing access to the Amarel cluster and associated research computing resources that have contributed to the results reported here. We also acknowledge the Cornell National Drosophila Species Stock Center and the laboratory of Daniel Matute for providing fly stocks used in this study.

