## Supplemental Figures for "Convergence and conflict among telomere specialized transposons across 60 million years of Drosophilid evolution"

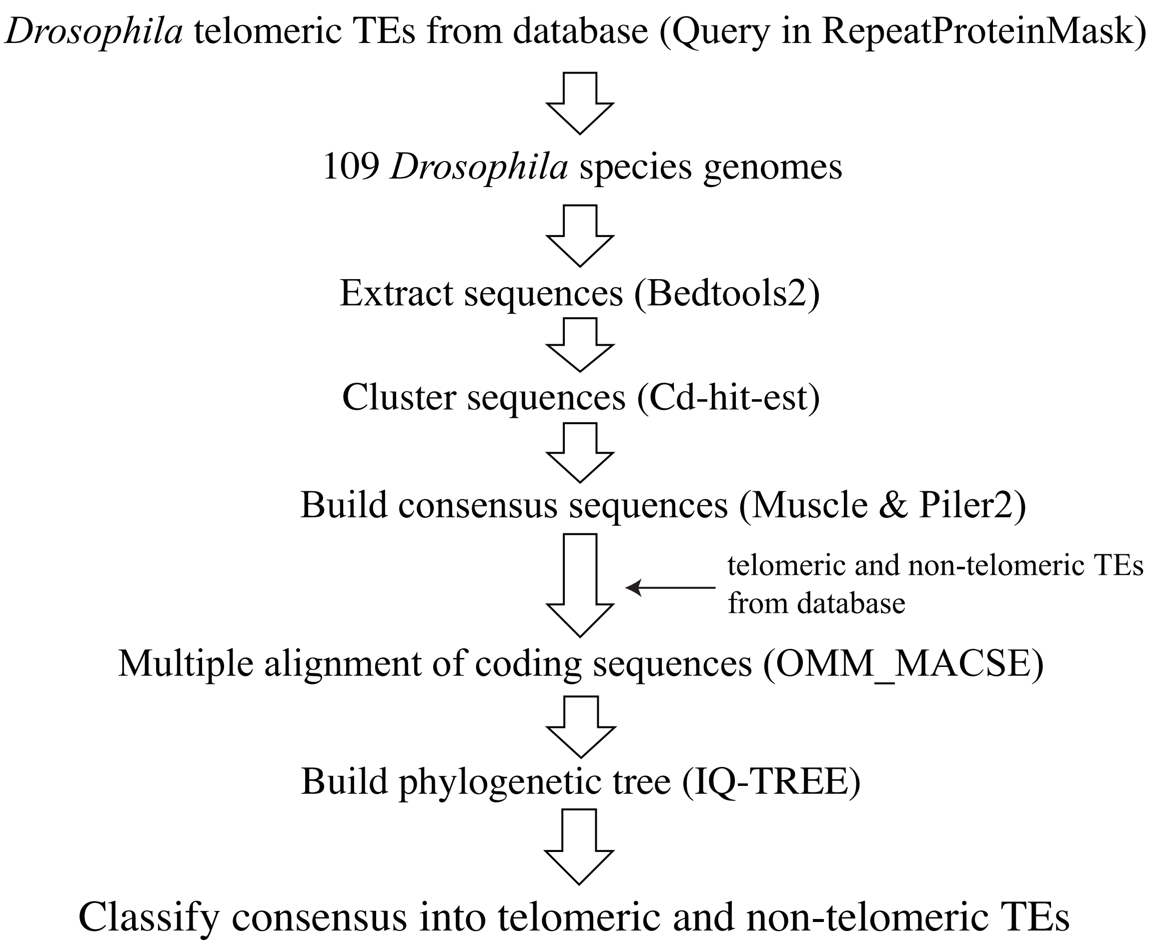


**Figure S1. Pipeline to identify telomeric retrotransposons across the *Drosophila* genus**. Known *Drosophila* telomeric TE peptides from Repbase were used to run *RepeatProteinMask* against 109 *Drosophila* species' genomes. *RepeatProteinMask* hits were extracted using BEDTOOLS and clustered with CD-HT-EST. Each cluster of sequences was aligned using MUSCLE and consensus sequences for each cluster were generated using PILER. OMM_MACSE was used to generate frameshift-aware amino acid translations of the ORF1 and ORF2 genes which were then used to construct phylogenetic trees with IQ-TREE. The ORF1 and ORF2 gene trees were used to assign each TE as putatively telomeric or non-telomeric. Manual curation was then used to identify head-to-tail arrays of putative telomeric elements and generate a full-length consensus sequence of each TE family. See Methods for additional details.

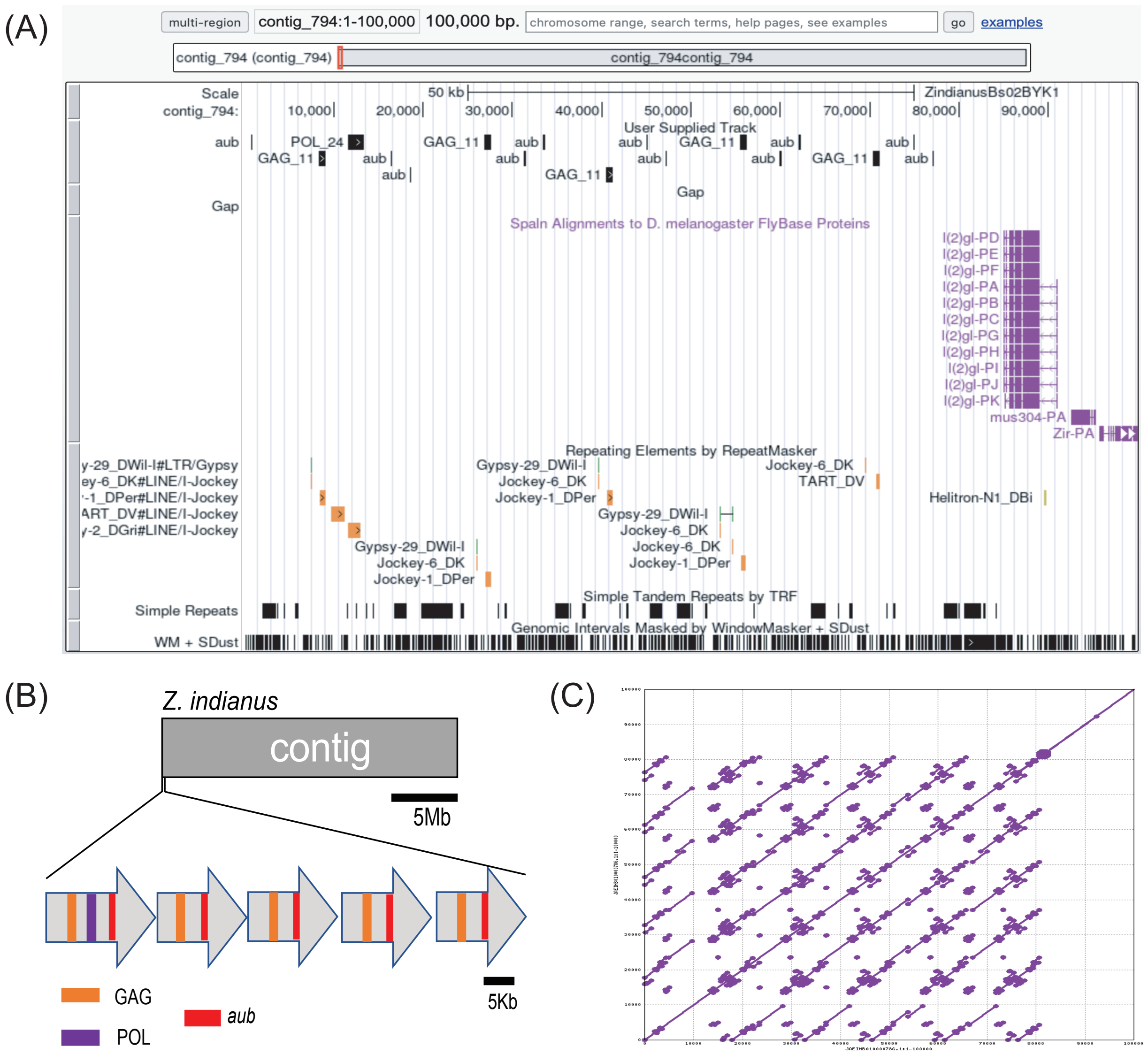


**Figure S2. Head-to-tail array of telomeric TEs on Muller Element B of *Z. indianus*.**
(A) UCSC Genome Browser custom track showing locations of RepeatMasker hits for ORF1 and ORF2 DNA queries (labeled as GAG_11 and POL_24) from putative *Z. indianus* telomeric TEs as well as BLASTN hits using *Z. indianus aubergine* mRNA as a query sequence. (B) A schematic representation of the head-to-tail array of telomeric TEs in the *Z. indianus* Muller B telomere. (C) A dot plot for the *Z. indianus* telomere region shown in (A).

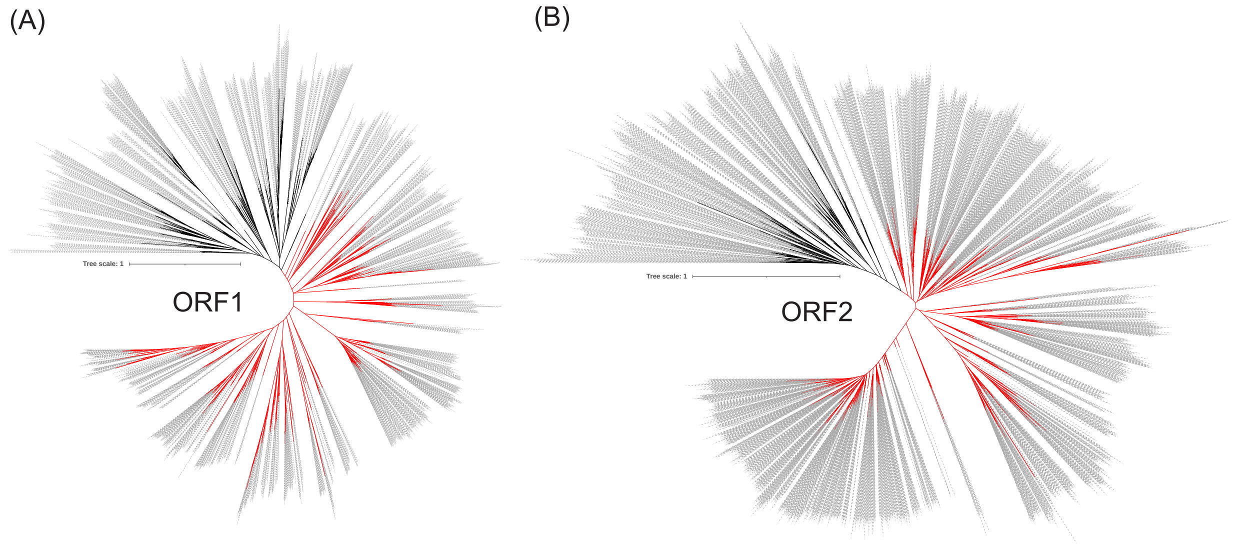


**Figure S3. ORF1 and ORF2 gene trees.** IQ-TREE gene trees of all ORF1 (A) and ORF2 (B) amino acid sequences identified from the pipeline. Each ORF was assigned as telomeric (black) or non-telomeric (red) based on whether it was more closely related to known telomeric or non-telomeric ORFs. Bootstrap supports for the telomeric clade are 89% and 76% for ORF1 and ORF2 trees, respectively.

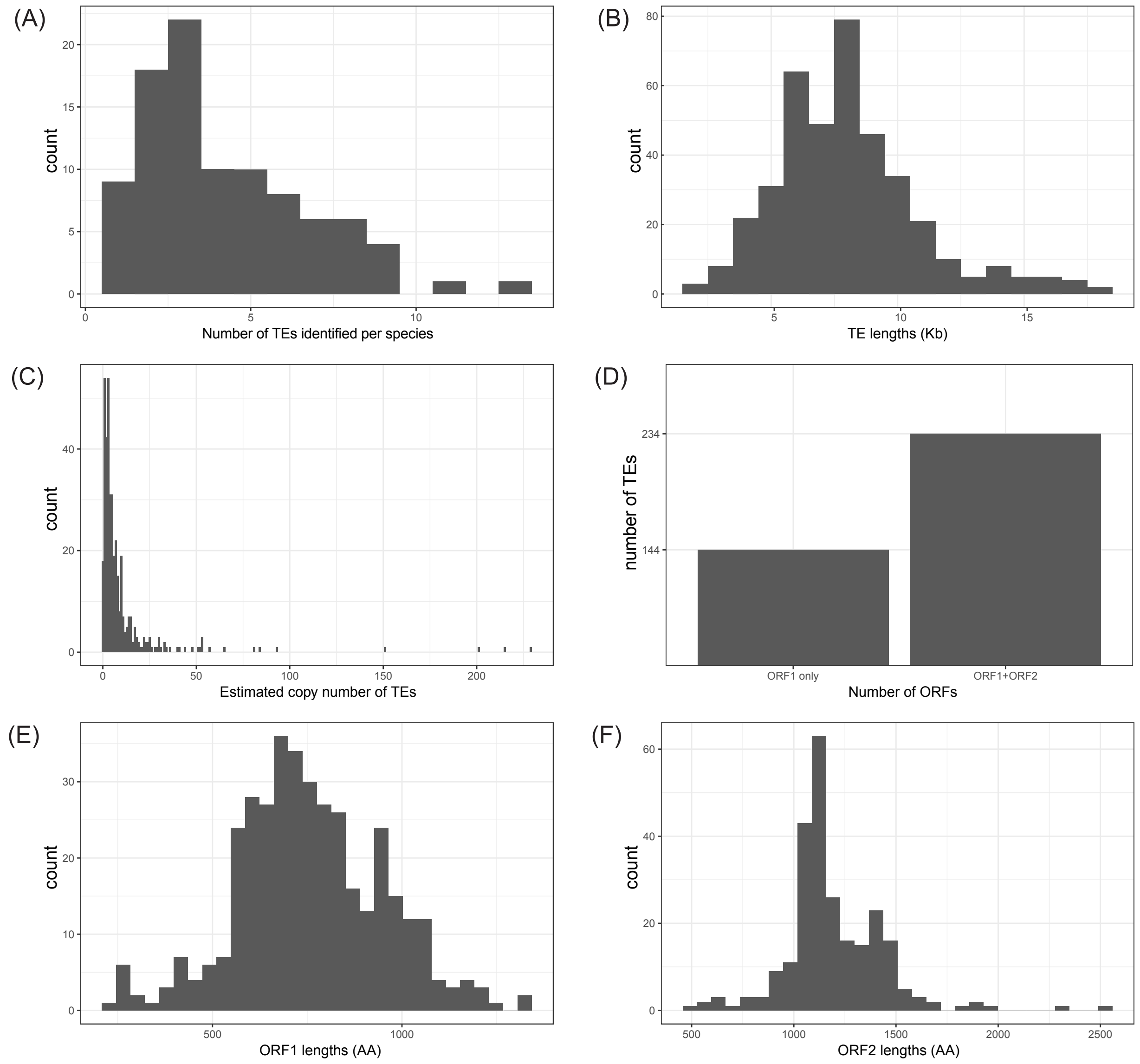


**Figure S4. Summary of telomeric TE properties across all species.** (A) The number of telomeric TEs identified per species. (B) DNA sequence length of full-length telomeric TE families. (C) Estimated genomic copy number of each telomeric TE family (D) Counts of telomeric TE families with ORF1 only versus those with both ORF1 and ORF2. (E & F) Amino acid sequence lengths of ORF1 and ORF2 from each telomeric TE.

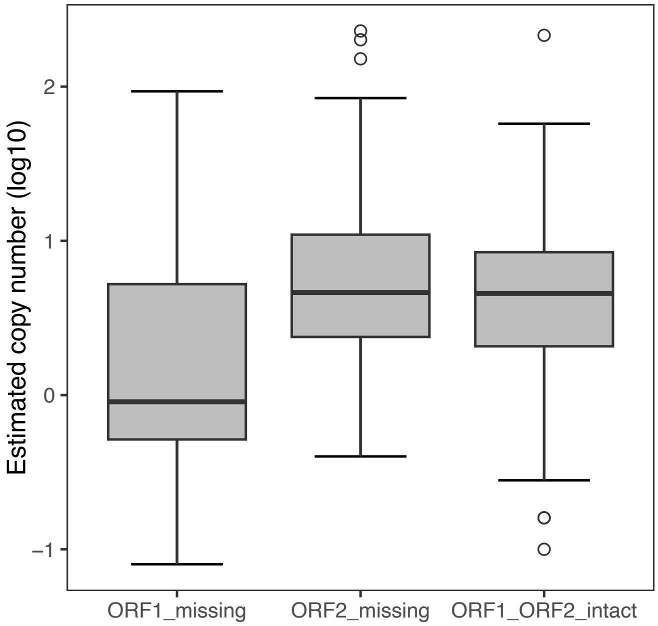


**Figure S5. TE copy number versus ORF contents.** We identified 18 TE families where ORF2 was present but ORF1 was missing. These TEs have significantly fewer genomic copies compared to both TE families where ORF2 is missing and TE families where both ORF1 and ORF2 are present, suggesting that the ORF1-negative TEs may be inactive, fragmented copies.

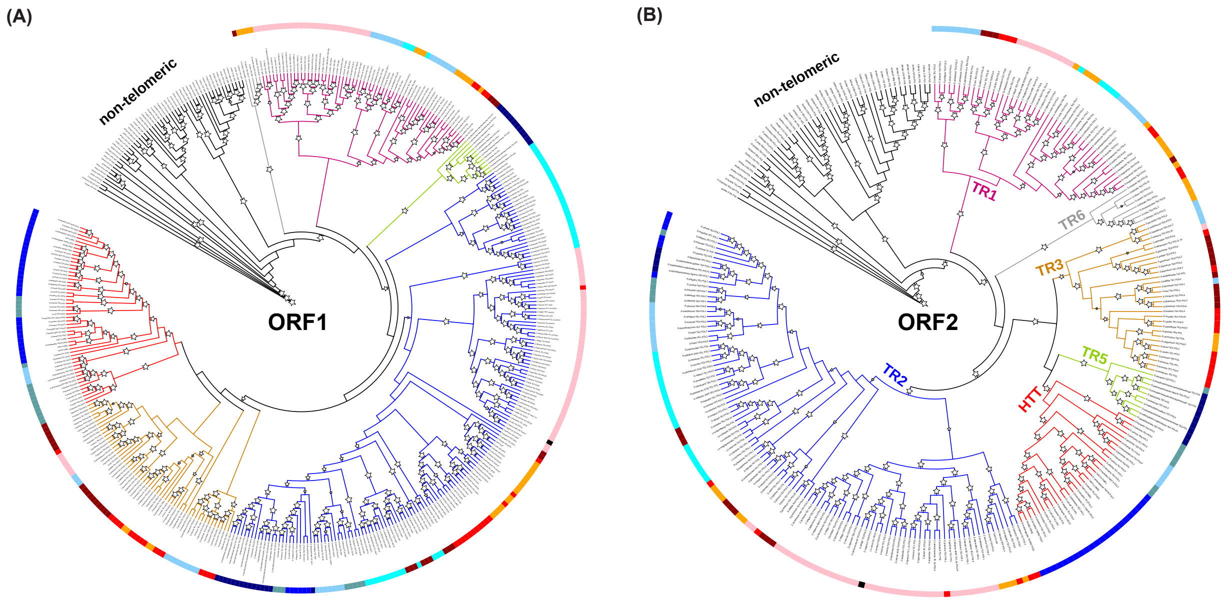


**Figure S6. Final telomeric TE gene trees**. ORF1 (A) and ORF2 (B) amino acid sequences from the manually curated telomeric TEs were aligned with ORFs from known non-telomeric TE families as outgroups (black branches) and trees were inferred using IQ-TREE (see Methods). The colored branches indicate major telomeric TE clades. The colors in the outer circle of the trees indicate major *Drosophila* species clades, as delineated in (Suvorov et al. 2022) (see Figure 1). Stars at each node indicate bootstrap support >= 85%.

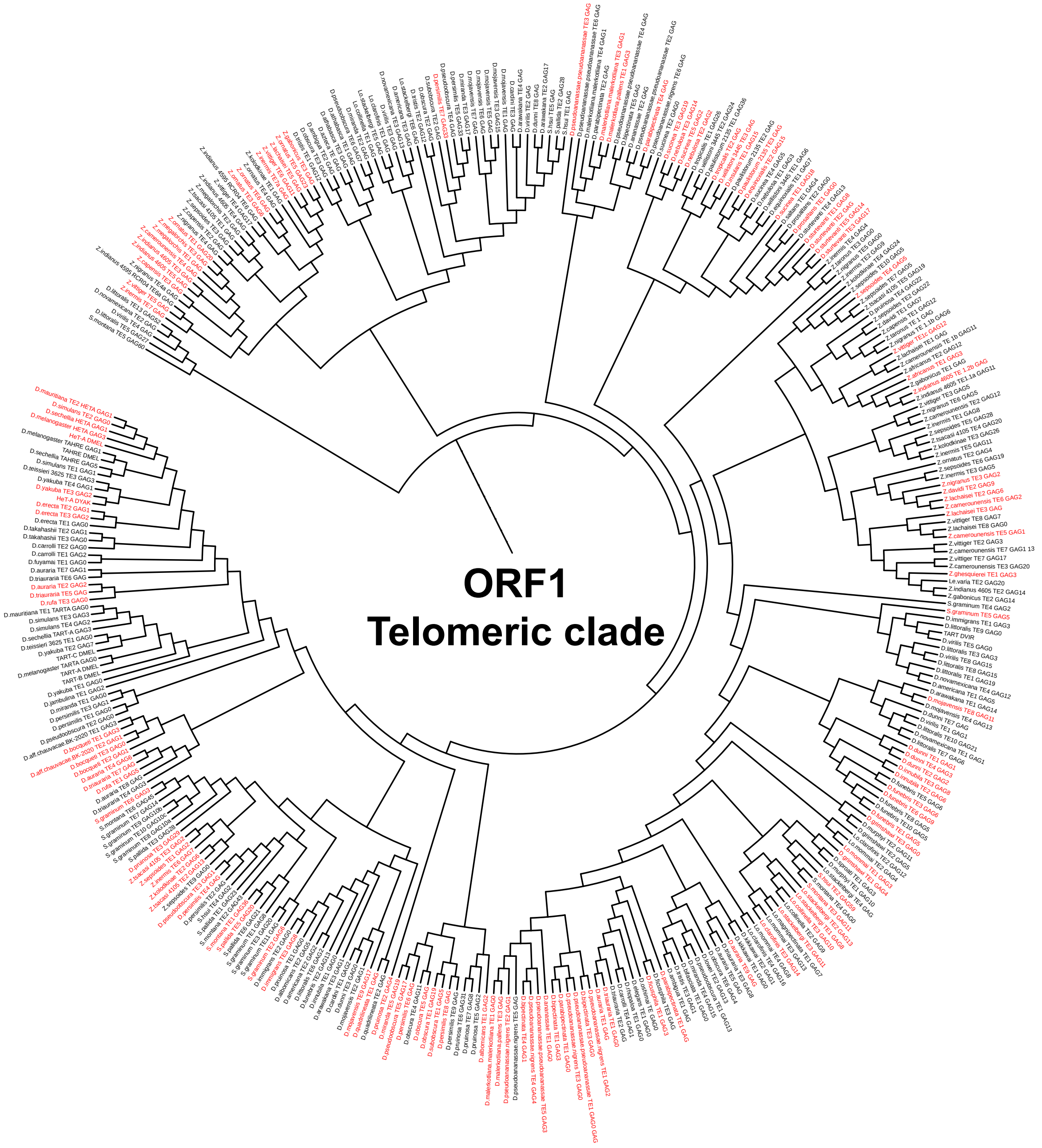


**Figure S7**. **Repeated loss of ORF2 from telomeric TEs.** ORF1-only telomeric TEs, which have lost ORF2, are shown in red. The largest monophyletic clade of ORF1-only TEs contains six elements. The presence of ORF1-clades suggests that new ORF1-only TE lineages can be derived from ancestral ORF1-only TEs. However, these lineages are short-lived, as indicated by the absence of large clades of ORF1-only TEs.

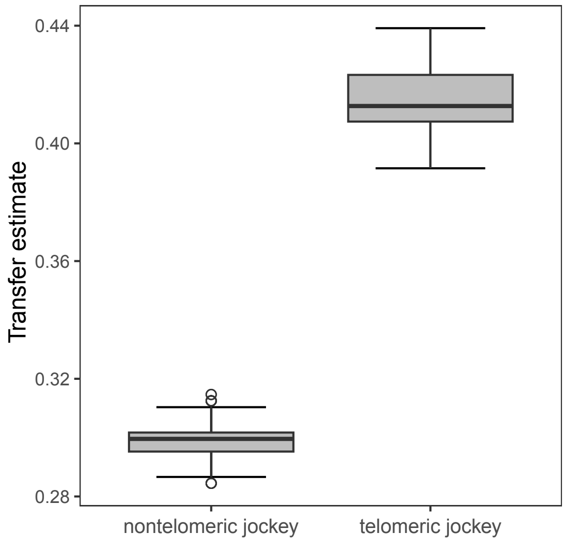


**Figure S8. Rates of horizontal transfer for non-telomeric versus telomeric Jockey clade TEs**. Horizontal transfer events were inferred using rangerDTL. The total number of transfer events for each group (i.e. telomeric versus non-telomeric) was then normalized by the total number of nodes in the corresponding tree. Boxplots show the distribution of these estimates across 1000 runs of rangerDTL.

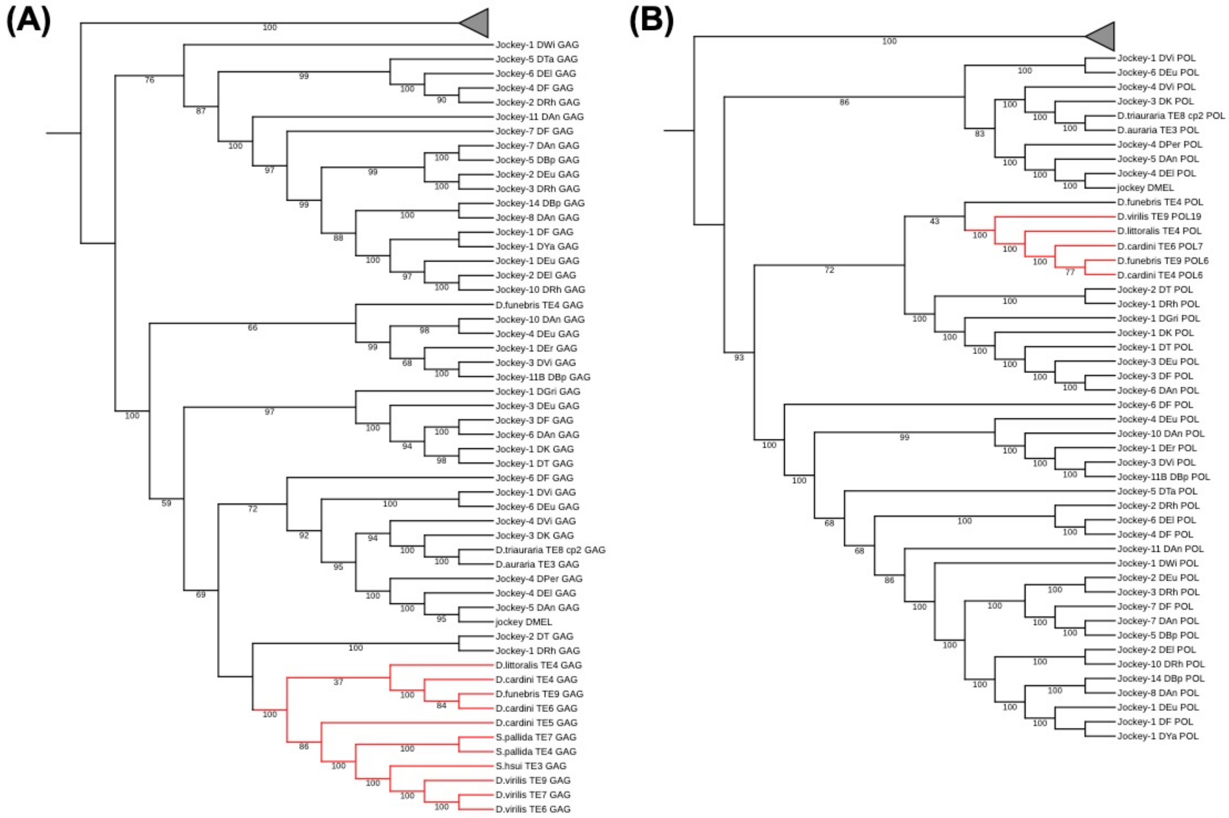


**Figure S9. Non-telomeric clade TEs at telomeres (NTTs).** Gene trees from ORF1 (A) and ORF2 (B) amino acid sequences. All known non-telomeric Jockey clade TEs from Repbase were included in addition to the NTT TEs identified here. The telomeric TE clade (grey triangle) has been collapsed. The NTT TEs (red branches) form a well-supported monophyletic subclade within the larger non-telomeric clade.

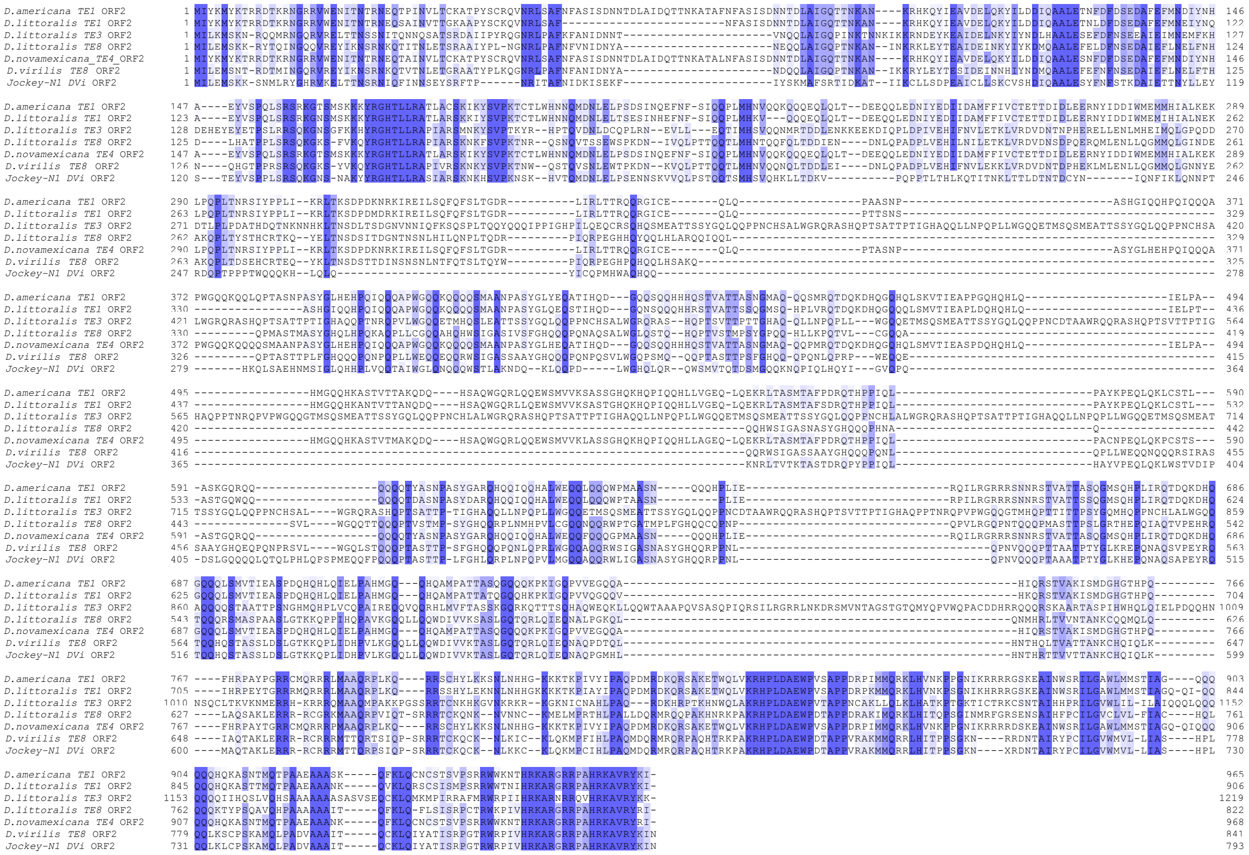

**Figure S10. Potential neofunctionalization of ORF2.** Seven telomeric TE families from members of the *virilis* species group (see Figure 3E) contain an ORF2 that shows no significant homology (via peptide BLAST) to the canonical ORF2 of other telomeric TE families. This unusual ORF2 also lacks both the endonuclease and reverse transcriptase conserved domains. Shown here is a multiple sequence alignment of the peptide sequences of this ORF2 from all seven TE families. Columns are shaded based on their level of conservation.

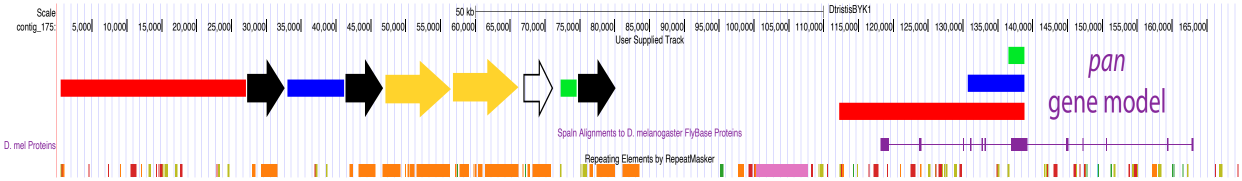


**Figure S11. *Pangolin* gene fragments in the dot chromosome telomere of *D. tristis*.** *D. tristis* UCSC Genome Browser screenshot showing the telomeric end of the dot chromosome. In this species, the protein-coding gene *pangolin* (*pan*) is located directly adjacent to the telomere while duplicated fragments of *pan* are found within the array of telomeric transposons. The duplicated *pangolin* fragments are quite large (>20 Kb in one case. We therefore conclude that the presence of the duplicated *pangolin* fragments in the *D. tristis* telomere may be due to unequal crossing over. In all of the other gene capture instances, the captured host gene (i.e. *nxf2, aub, piwi, or CG12520*) is at least 1.7 Mb from the telomere, making unequal crossing over unlikely in these instances. Closed black and yellow arrows represent full-length copies of the two telomeric TE families found at this telomere. The open black arrow is a partial-length copy, missing ORF2. The colored rectangles show the three *pan* duplications within the telomere as well as their parental copy within the *pan* locus.

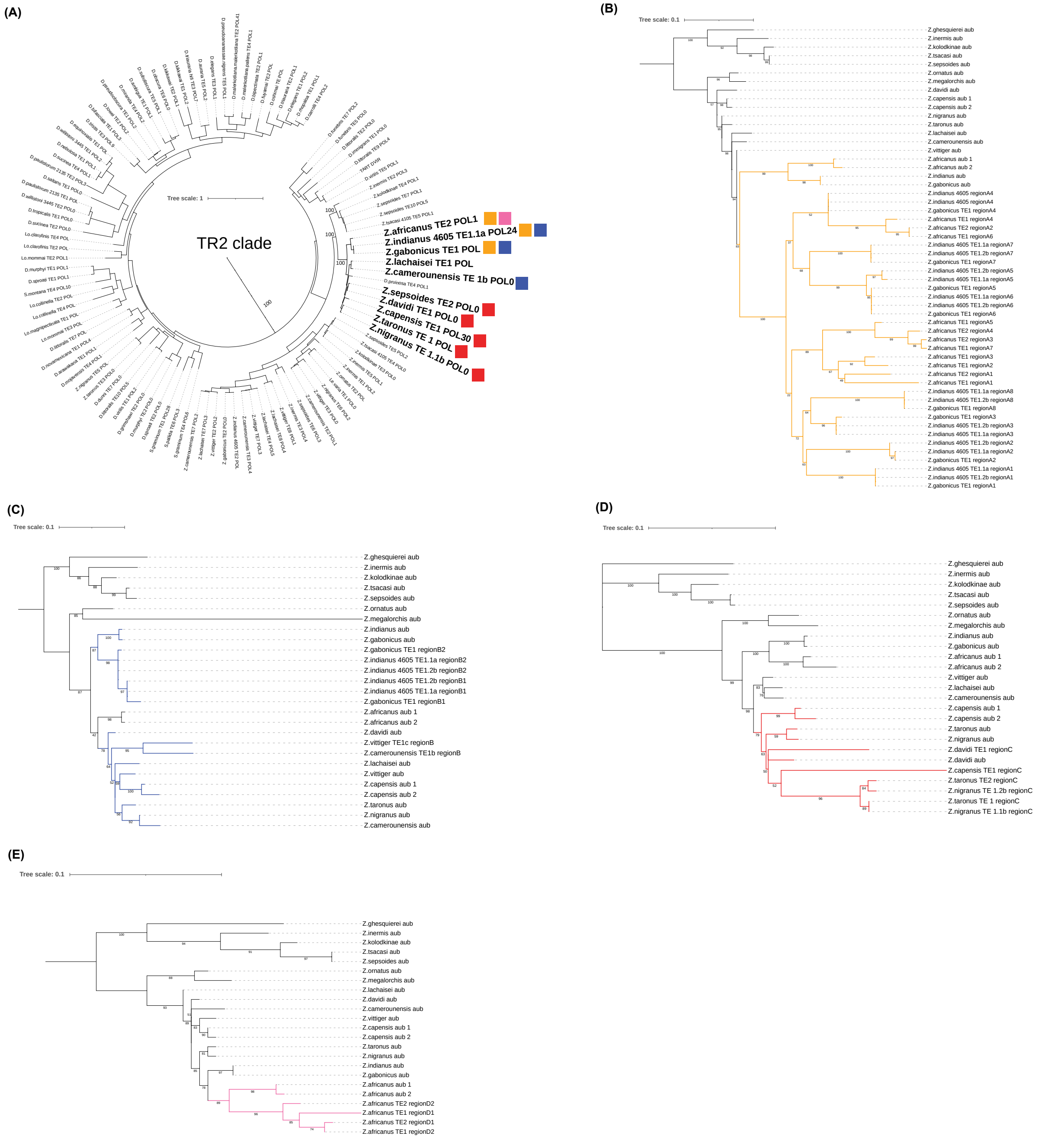


**Figure S12. Phylogenetic reconstruction of gene capture events in *Zaprionus*.** (A) With the exception of *D. pruinosa* TE4 (which does not contain an *aub* fragment), the *Zaprionus* telomeric TEs that captured *aub* sequences (bold text) form a monophyletic clade in the ORF2 gene tree. The colored squares show which region of *aub* was captured by each TE family (see Figure 5A). (B – E) To infer the most recent common ancestor of the *aub*-like TE sequences and their host gene progenitors, we created separate gene trees for each captured region of *aub*. Each tree was created from each region of the *aub* gene that was captured by at least one TE family (i.e. Regions A-D, see Figure 5A) along with the corresponding *aub*-like TE sequences. (B) Region A: all *aub*-like TE sequences form a single clade, consistent with a single capture event for this region. This TE clade is most closely related to the *aub* gene sequences from *Z. africanus*, *Z. indianus*, and *Z. gabonicus*, suggesting that the Region A gene capture event occurred in the common ancestor of these three species. (C) Region B: the *aub*-like TE sequences form two separate clades, consistent with two independent capture events for this region. Event 1: This TE clade is most closely related to the *aub* gene sequences from *Z. indianus* and *Z. gabonicus*, suggesting that this capture event occurred in the common ancestor of these two species. Event 2: The *Z. vittiger* and *Z. camerounensis aub*-like TE sequences are most closely related to the *aub* gene sequences from *Z. camerounensis*, *Z. lachaisei*, *Z. vittiger*, *Z. davidi*, *Z. taronus*, and *Z. capensis*, suggesting that this capture event occurred in the common ancestor of these six species, followed by two independent losses of the captured sequence, one in the ancestor of *Z. davidi*, *Z. taronus*, and *Z. capensis* and one in *Z. lachaisei*. (D) Region C: all *aub*-like TE sequences form a single clade, consistent with a single capture event for this region. This TE clade is most closely related to the *aub* gene sequences from *Z. davidi*, *Z. taronus*, and *Z. capensis* (note: *Z. nigranus aub* is present in this clade due to horizontal transfer (or introgression) of *aub* from *Z. taronus* to *Z. nigranus*, see Figure S15), suggesting that the Region C gene capture event occurred in the common ancestor of these three species. (E) Region D: the *Z. africanus aub*-like TE sequences form a single clade, sister to the *Z. africanus aub* gene sequences, suggesting that this capture event occurred specifically in the *Z. africanus* lineage.

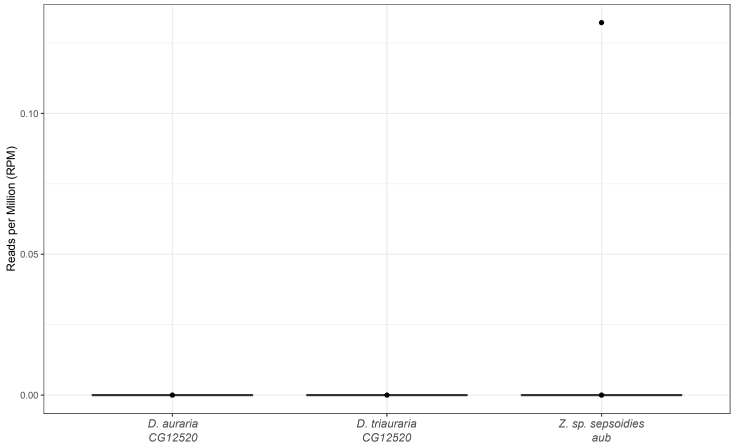


**Figure S13. Two older gene capture fragments do not appear to be capable of inducing piRNA production from their host genes.** A telomeric TE family in *D. auraria* and *D. triauraria* captured a fragment of the host gene *CG12520*. The identity between the captured fragment and host gene is ~79%. Similarly, the identity between *Z. sp sepsoides aub* and the *aub*-like sequence carried by its telomeric TE is ~82%. The box plots above show normalized piRNA abundance (calculated in 80 bp sliding windows across the gene transcripts).

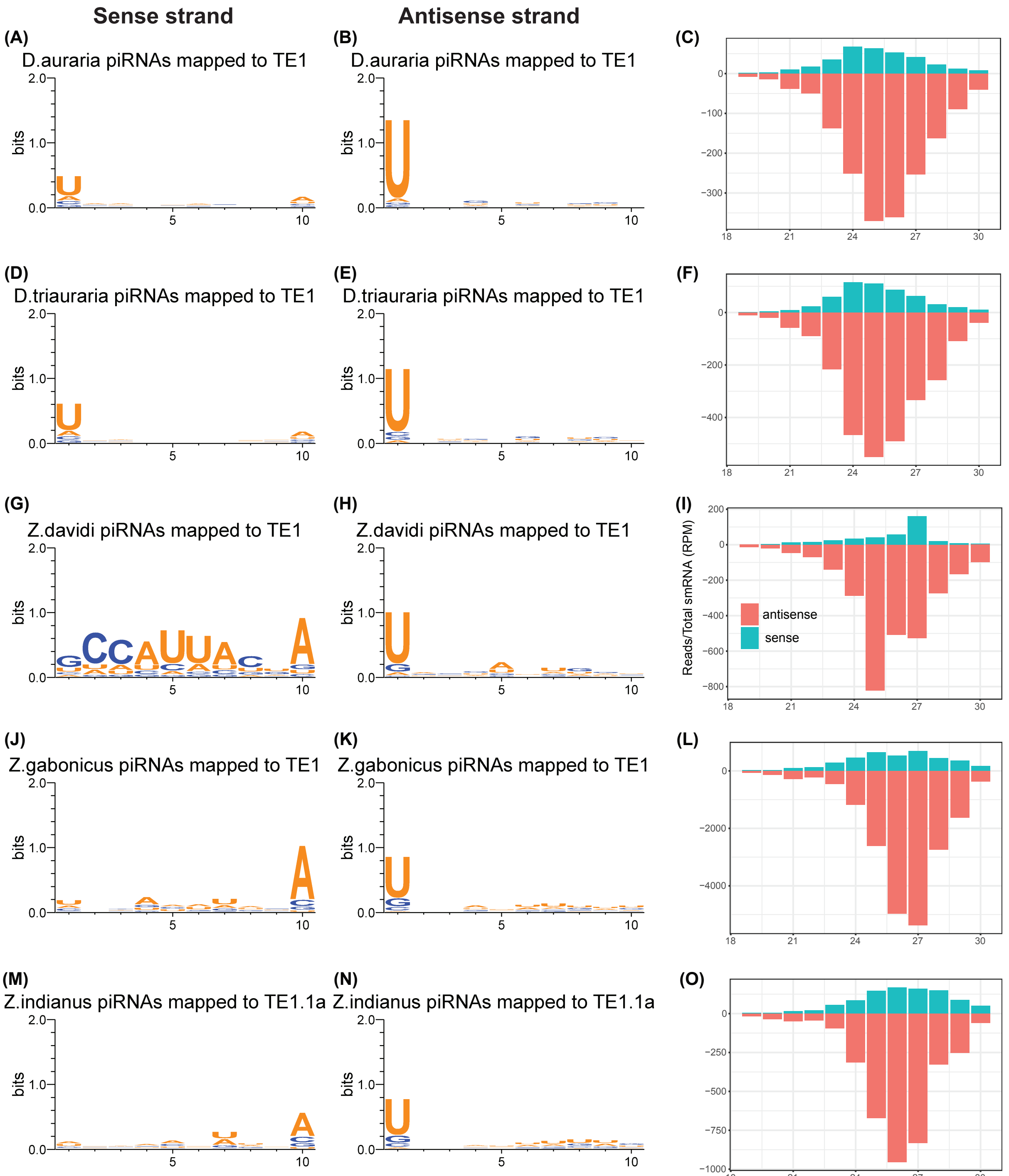


**Figure S14. Telomeric TE-derived piRNAs.** Nucleotide frequency sequence logos made from 10 bp 5' ends of piRNAs mapped to host gene capture TEs in the sense and antisense orientations along with the length distribution and abundance of all small RNAs aligned to the same TEs.

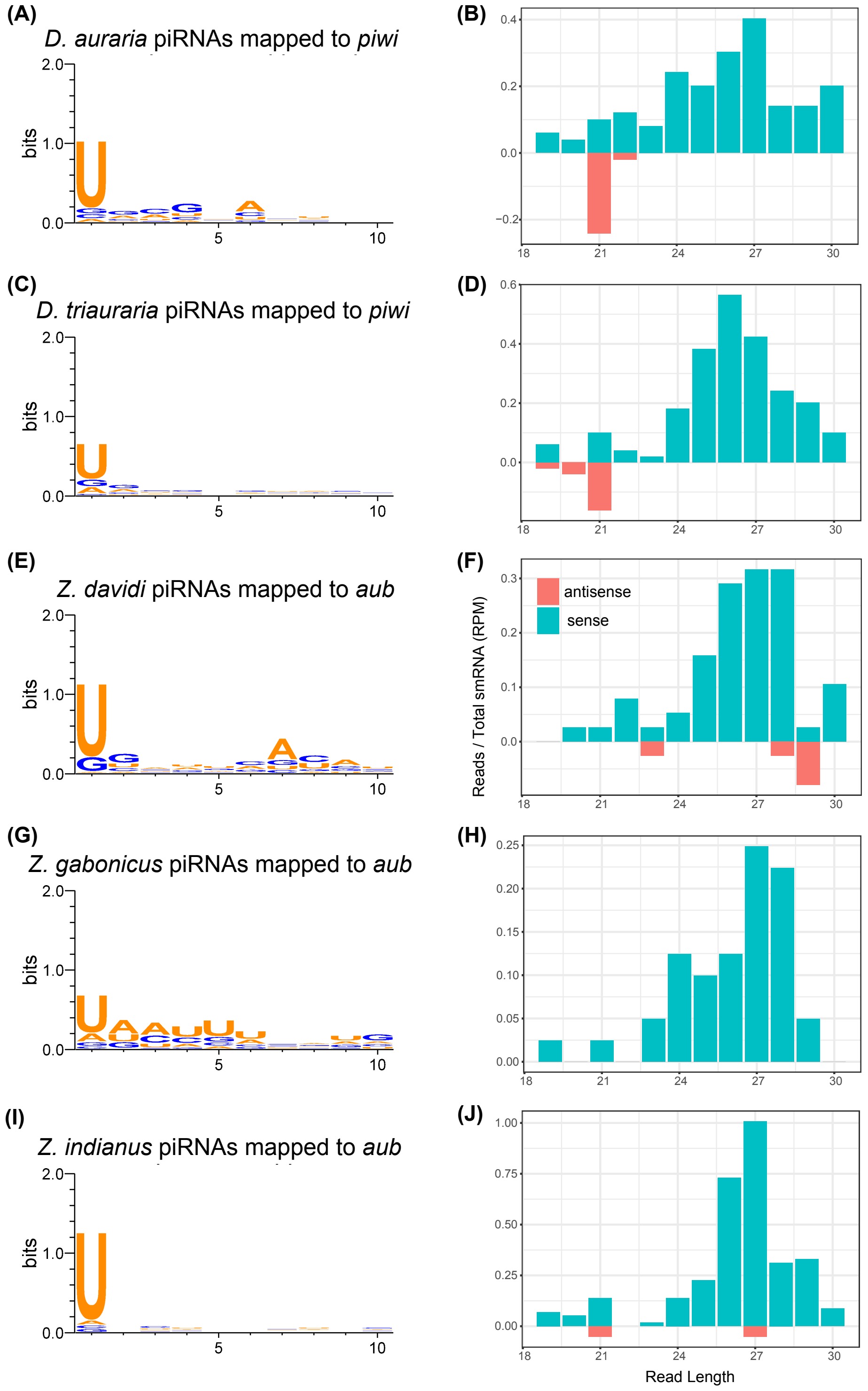


**Figure S15. Host gene-derived piRNAs.** Nucleotide frequency sequence logos made from 10 bp 5' ends of piRNAs mapped to the sense strand of the host gene that was captured by a telomeric TE in (i.e. *piwi* or *aub*) along with the length distribution and abundance of all small RNAs aligned to the same gene. Note that antisense sequence logos are not shown due to the lack of antisense piRNAs.

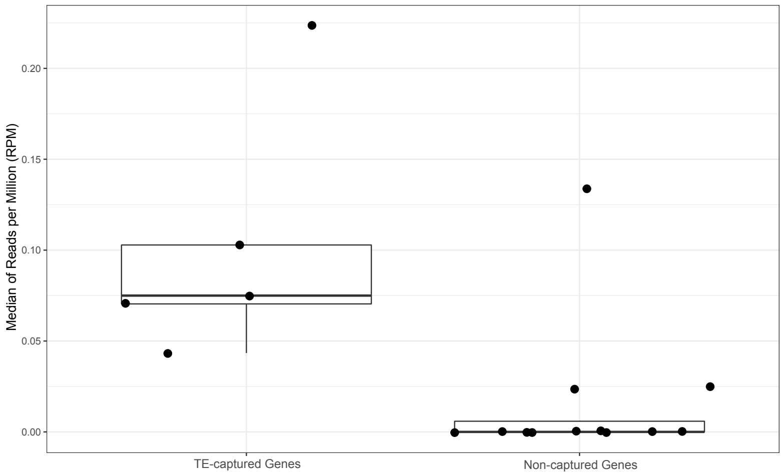


**Figure S16. PiRNA abundance from captured versus control host genes.** Median normalized piRNA abundance from the host genes captured by telomeric TEs (*piwi* for *D. auraria* and *D. triauraria*, and *aub* for *Z. davidi*, *Z. gabonicus*, and *Z. indianus*) versus the control host genes not captured by telomeric TEs (*aub* for *D. auraria* and *D. triauraria, Z. quesquierei*, *Z. kolodkinae*, *Z. sp. megalorchis*, and *Z. sp. sepsoides,* and *piwi* for *Z. davidi*, *Z. gabonicus*, *Z. indianus, Z. quesquierei*, *Z. kolodkinae*, *Z. sp. megalorchis*, and *Z. sp. sepsoides*), calculated in 80 bp sliding windows across the gene transcript. The captured genes produce significantly more piRNAs compared to their controls (Wilcoxon test: $P$ = 0.0036).

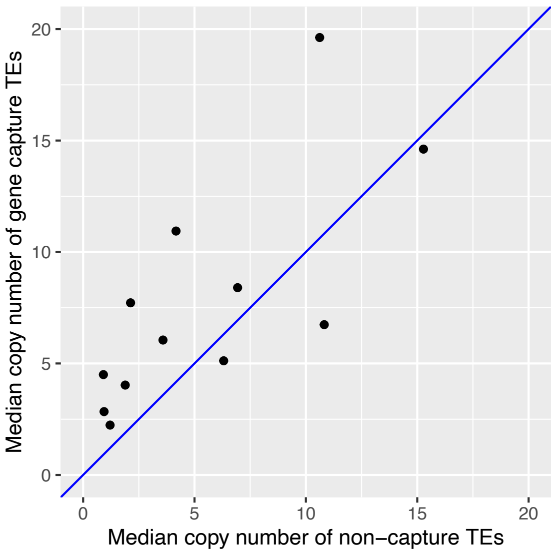


**Figure S17. Gene capture telomeric TEs tend to be present at higher copy numbers compared to their non-capture counterparts.** Each dot represents a single species whose genome contains both gene-capture and non-capture telomeric TEs. The telomeric TE families that have captured a host gene fragment have significantly higher copy numbers compared to the other non-capture telomeric TEs in the same genome (paired Wilcoxon test, one-sided $P$ = 0.02).

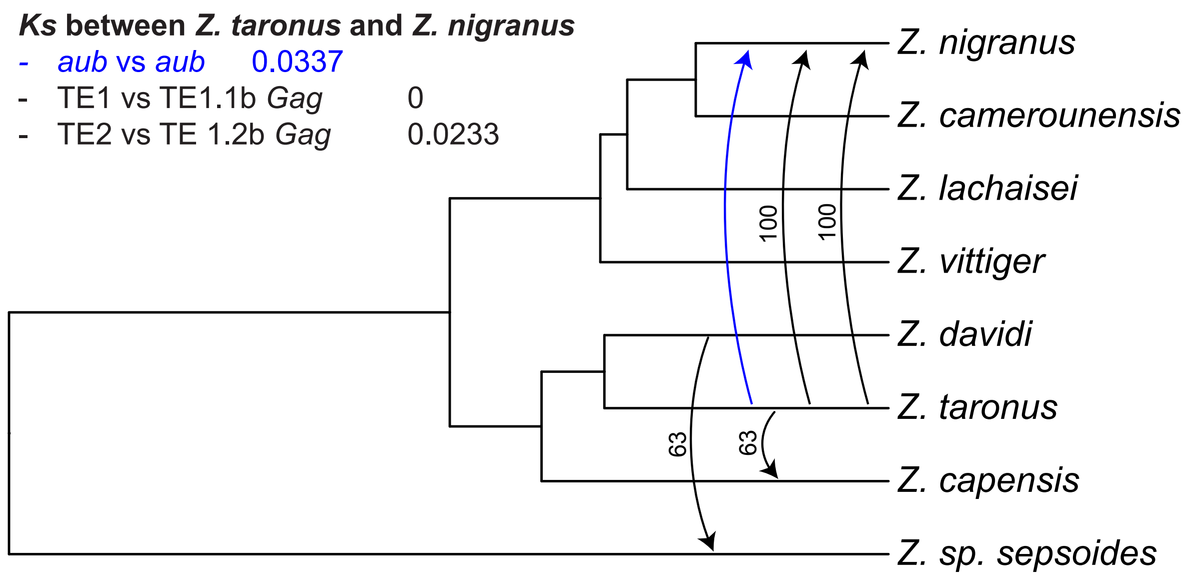


**Figure S18. TE tree/species tree reconciliation from *Zaprionus* gene capture TEs.** Gene tree/species tree reconciliation identified four possible horizontal transfer events involving *Zaprionus* gene capture TEs (black arrows, numbers indicate percent of reconciliation solutions supporting each event). Also shown is the horizontal transfer/introgression of *aub* from *Z. taronus* to *Z. nigranus* (blue arrow). The relative timing of the *aub* versus TE transfers from *Z. taronus* to *Z. nigranus* was inferred based on Ks values shown above.

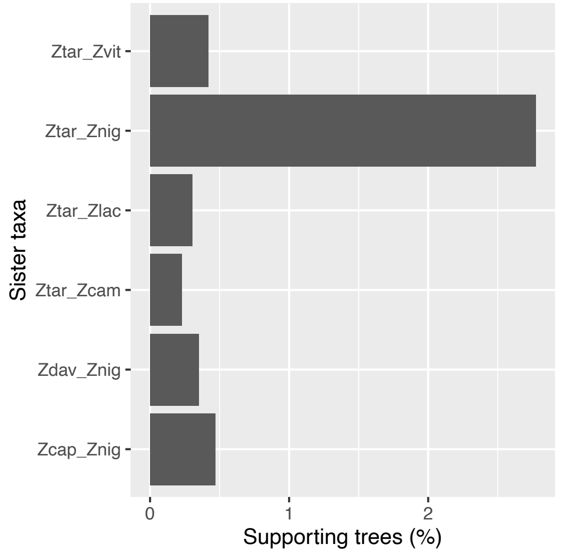


**Figure S19**. **Evidence of gene flow between *Z. taronus* and *Z. nigranus*.** We analyzed 2,500 single-copy ortholog gene trees from (Suvorov et al. 2022). We quantified the number of gene trees showing each sister taxa relationship above and found a significant excess of trees supporting *Z. taronus* and *Z. nigranus* as sister taxa, consistent with introgression and/or gene flow between these two species (binomial test $P < 2.2e-16$).
